# What are oxytocin assays measuring? Epitope mapping, metabolites, and comparisons of wildtype & knockout mouse urine

**DOI:** 10.1101/2022.03.03.482682

**Authors:** Gitanjali E. Gnanadesikan, Elizabeth A.D. Hammock, Stacey R. Tecot, Rebecca J. Lewis, Russ Hart, C. Sue Carter, Evan L. MacLean

## Abstract

Oxytocin has become a popular analyte in behavioral endocrinology in recent years, due in part to its roles in social behavior, stress physiology, and cognition. Urine samples have the advantage of being non-invasive and minimally disruptive to collect, allowing for oxytocin measurements even in some wild populations. However, methods for urinary oxytocin immunoassay have not been sufficiently optimized and rigorously assessed for their potential limitations. Using samples from oxytocin knockout (KO) and wildtype (WT) mice, we find evidence of considerable interference in unextracted urine samples, with similar distributions of measured oxytocin in both genotypes. Importantly, although this interference can be reduced by a reversed-phase solid-phase extraction (SPE), this common approach is not sufficient for eliminating false-positive signal on three immunoassay kits. To better understand the source of the observed interference, we conducted epitope mapping of the Arbor Assays antibody and assessed its cross-reactivity with known, biologically active fragments of oxytocin. We found considerable cross-reactivity (0.5-52% by-molarity) for three fragments of oxytocin that share the core epitope, with more cross-reactivity for longer fragments. Given the presence of some cross-reactivity for even the tripeptide MIF-1, it is likely that many small protein metabolites might be sufficiently similar to the epitope that at high concentrations they interfere with immunoassays. We present a new mixed-mode cation-exchange SPE method that minimizes interference—with knockout samples measuring below the assay’s limit of detection—while effectively retaining oxytocin from the urine of wildtype mice. This method demonstrates good parallelism and spike recovery across multiple species (mice, dogs, sifakas, humans). Our results suggest that immunoassays of urine samples may be particularly susceptible to interference, even when using common extraction protocols, but that this interference can be successfully managed using a novel mixed-mode cation exchange extraction.

## 1 Introduction

Oxytocin has been the focus of considerable research effort over the last several decades, especially in behavioral endocrinology and pharmacology. Early work demonstrated its importance for maternal behavior (Pederson & Prange, 1979; Van Leengoed et al., 1987) and pair-bonding (reviewed in Carter et al., 1995). Recently, evidence has been mounting for oxytocin’s role in a myriad of social behaviors and cognitive processes, as well as stress physiology (reviewed in Feldman, 2012; Carter, 2014; Froemke & Young, 2021).

Making sense of the oxytocin literature can be challenging, partially due to the variety of experimental methods used. For example, administration of exogenous oxytocin can produce dosage-dependent results (Bales et al., 2007; Cardoso et al., 2013; Peters et al., 2014), and the effects of administration are not necessarily the same as the effects of endogenous oxytocin (Crockford et al., 2014). While administration studies may be fruitful for pharmacology, studies of endogenous oxytocin are crucial for understanding the biology of the oxytocin system, its function, and its evolution. However, studies of endogenous oxytocin concentrations have often struggled to reliably measure oxytocin, and there are disagreements regarding the best approaches for measuring oxytocin, especially by immunoassay (e.g. McCullough et al., 2013; Lefevre et al., 2017; MacLean et al., 2019). Furthermore, endogenous oxytocin measurements and patterns of change are often uncorrelated across biological matrices, especially urine (Amico et al., 1987; Feldman et al., 2011; Weber et al., 2017). Complicating these comparisons, sample preparation techniques—such as extraction—that have been developed and validated in one matrix are often assumed to work equally well in other matrices, despite their different compositions.

Using blood plasma samples from oxytocin knockout (KO) and wildtype (WT) mice, we previously demonstrated that while some commercially available immunoassay kits (i.e. Enzo Life Sciences) are susceptible to interference, others (i.e. Arbor Assays) are not, even with unextracted (diluted 1:8) mouse plasma samples (Gnanadesikan et al., 2021). Here, we turn this approach towards improving and validating urinary oxytocin (uOT) measurements. While plasma samples are often available in laboratory research settings, blood collection is often impossible or disruptive. In studies of wild populations, blood collection usually involves trapping animals, and even in captive settings or human studies, it may affect behaviors of interest. Thus, the advantages of non-invasive hormone measures for behavioral endocrinology have long been appreciated both in general (Strier & Ziegler, 2005) and for oxytocin research specifically (Crockford et al., 2014). However, developing reliable methods has been challenging.

Urine samples provide a promising avenue for non-invasive behavioral endocrinology for several reasons: 1) urine can be caught in a container, on an absorbent material, or pipetted up off surfaces without handling the animal; 2) urination may occur throughout the day, enabling collection at multiple time points; 3) many species produce volumes of urine that are sufficient to assay multiple analytes in the same sample.

### 1.1 Urinary oxytocin and behavior across species

Many studies have already utilized uOT measurements and found interesting associations with social behavior. In humans, uOT has been associated with perceptions of a romantic partner’s bonding behavior, regardless of gender (Algoe et al., 2017), and with performance on a facial visual search task in men (Saito et al., 2014). In children, a mother’s touch and speech after a stressful event increased uOT while decreasing salivary cortisol (Seltzer et al., 2010). Interestingly, in mothers, uOT was higher after interacting with other people’s children compared to their own children, perhaps due to oxytocin’s role in the onset—rather than maintenance—of maternal behavior, or perhaps simply due to a novelty effect (Bick & Dozier, 2010).

Increasingly, uOT has been used in studies of human-animal interactions. Nagasawa et al. (2009, 2015) demonstrated that uOT in dogs and their owners was associated with mutual gazing. Mitsui et al. (2011) suggested that oxytocin could be used as a biomarker of positive emotion, finding that dogs’ uOT increased after feeding, exercising, and petting, but not drinking water. In contrast, Marshall-Pescini et al. (2019) found that affiliative physical contact between dogs and people did not produce consistent changes in uOT levels measured in either the dogs or their owners. Wirobski, Range, et al. (2021) found that in pet dogs, uOT was associated with physical contact with their owner; however, in hand-raised but pack-living dogs and wolves, there was no association between measured uOT and physical contact with a bonded human, suggesting the importance of the animal’s life experience and the strength of the human-animal bond.

Primatologists have also made use of uOT measurements in a variety of lab and field studies. Using SPE and high-performance liquid chromatography (HPLC), Seltzer & Ziegler (2007) found that a captive male common marmoset’s uOT was lower during a 48-hour social isolation period than during the subsequent visual reintroduction to his mate. In captive tamarins, Snowdon et al. (2010) found high levels of individual uOT variation, with pair-bonded mates exhibiting similar concentrations that were partially explained by their affiliative behaviors. Similarly, in captive common marmosets, strongly bonded pairs exhibited synchronous fluctuations in uOT concentrations (Finkenwirth et al., 2015), and uOT was associated with infant care behaviors such as infant-licking and proactive food sharing in all group members (Finkenwirth et al., 2016). In field studies of chimpanzees, uOT has been associated with social bonding (Crockford et al., 2013), food sharing (Wittig et al., 2014), and intergroup conflict (Samuni et al., 2017). In baboons, higher uOT concentrations have been associated with a female’s estrous stage and their proximity maintenance with consort partners (Moscovice & Ziegler, 2012), as well as female-female sexual behavior (Moscovice et al., 2019).

Thus, uOT studies have already documented a variety of intriguing associations with social behavior, and there is some consistency—but also many differences—across studies and species.

### 1.2 Challenges for immunoassay of urinary neuropeptides

However, the specificity and reliability of uOT techniques has not always been tested rigorously. Even when extensive validation work has been conducted (e.g. Seltzer et al., 2010; Wirobski, Schaebs, et al., 2021), assessment of false-positive signal has not been possible, lacking a “ground truth”. Additionally, although many studies have employed some form of extraction, it remains unknown to what extent these methods are sufficiently selective to retain oxytocin while eliminating potentially interfering molecules. Urine contains a large quantity and variety of metabolites (Bouatra et al., 2013) that we expect might interfere with immunoassays, and thus extraction and immunoassay specificity may be particularly important for this matrix. In this study, we therefore tested multiple methods on both oxytocin KO and WT mice, searching for techniques that minimized interference in the KO samples while maximizing signal in the WT samples.

The specificity of immunoassays can be affected by many aspects of their chemistry. The antibody is particularly important, as different antibodies may bind different portions of a molecule. This region of an antigen that is bound by the antibody is known as the epitope. Aiming to shed light on potential sources of interference, we worked with Pepscan Presto BV (Lelystad, The Netherlands) to perform epitope mapping of the Arbor Assays antibody. Motivated by these results, we also tested the cross-reactivity of the Arbor Assays antibody with three known, biologically active fragments of oxytocin (Uvnäs Moberg et al., 2019), as well as the effects of two extraction methods on retention of these fragments.

Lastly, using a mixed-cation exchange method that minimized interference in knockout mouse samples, we tested its analytical validity across a variety of species (mice, sifakas, dogs, and humans).

## 2 Methods

### 2.1 Mouse subjects and sample collection

As in our previous work (Gnanadesikan et al., 2021), *Oxt*^*tm*1*Zuk*^ mice (Nishimori et al., 1996) were maintained on a C57BL/6J background and bred at Florida State University; all breeding and handling procedures were approved by the Institutional Animal Care and Use Committee (IACUC) following the Guide for the Care and Use of Laboratory Animals (Protocols 2020-00021 and 2020-00042). *Oxt*^+*/*−^ breeder pairs were continuously housed. The first morning of the appearance of a litter was noted as postnatal day 0 (P0). Mice were weaned, tagged, and tailed for genotyping on P21 and group housed by sex. After genotyping, same-sex mice were re-housed by genotype, so that only mice of the same genotype were housed together. Mice were housed on a 12:12L:D cycle in open wire-top caging with wood chip bedding and provided ad libitum food (LabDiet Rodent 5001) and water. In total, samples from 159 mice of this strain (WT: 73, KO: 86) were used in this study. Additionally, adult C57BL/6J (B6) wildtype mice were purchased from The Jackson Laboratory (total n = 20).

Urine samples were passively collected from adult male and female mice during routine handling on the day of planned euthanasia for other purposes. During live handling to confirm the mouse identification (ID number) by ear tag prior to euthanasia, the mouse was held over a microcentrifuge tube to collect urine. After confirming the mouse ID number, the tube was closed, labeled with the ID number, and frozen. Samples were shipped frozen to the University of Arizona on dry ice and stored at -80 °C until time of assay. Each urine sample, identifiable by ID number, was assigned a genotype code by E.H. to enable experimental design, assay layout planning, and sample pooling (within genotype when necessary), while retaining sample blinding for G.E.G. for assays directly comparing knockout and wildtype individuals.

#### 2.1.1 Genotyping

DNA from tail samples was genotyped for *Oxt* alleles using a common forward primer (5’-TCAGAGATTGAACAAGACGCC) and specific reverse primers for WT (5’-TCAGAGCCAGTAAGCCAAGC) and KO (5’-ACTTGTGTAGCGCCAAGTGC). Using a hot start and 40 cycles of 30 s each of 94 °C, 57 °C, and 72 °C, the primers generated a wildtype allele of approximately 500 bp and a knockout allele of approximately 180 bp, resolved by gel electrophoresis.

#### 2.1.2 Samples in each analysis

For a summary of mouse samples used in this study, see table A2. The first experiment was conducted on unextracted urine samples: 10 wildtype *Oxt*^+*/*+^ samples (7 female, 3 male) and 14 knockout *Oxt*^−*/*−^ samples (8 female, 6 male). The second experiment, as well as the methods development work reported in the appendix used a series of sample pools. In total, these pools used samples from 76 wildtype individuals (mix of C57BL/6J and *Oxt*^+*/*+^) and 62 knockout individuals (*Oxt*^−*/*−^). For experiment three, some individual samples were pooled across 2-3 individuals within the same—blinded—genotype code, due to limited sample volume. All samples were from male mice. Six individual *Oxt*^+*/*+^ wildtype samples were used; 10 individual *Oxt*^−*/*−^ knockout samples were pooled as necessary to create 6 pooled samples of 1-3 individuals each.

### 2.2 Sifaka samples

Verreaux’s sifaka (*Propithecus verreauxi*) samples were collected from a habituated wild population at Ankoatsifaka Research Station in Kirindy Mitea National Park, Madagascar, under IACUC Protocol #13-470 from the University of Arizona and IACUC Protocol #2017-00152 from the University of Texas at Austin. Fresh urine samples were collected from known, identified individuals by catching the stream directly with foil, catching the stream with foil off of tree trunks, or pipetting urine off of foliage with disposable pipettes. Samples were pipetted into cryovials, which were immediately placed into a thermos with either instant ice packs or tubes of frozen water until transferred into a liquid nitrogen tank within 4 hours. Samples were kept frozen in a liquid nitrogen dewar during transport from the field site to Antananarivo and while awaiting shipment to the US. They were shipped to the University of Arizona in a charged liquid nitrogen shipper and frozen at -80 °C until analysis at the Laboratory of the Evolutionary Endocrinology of Primates (LEEP).

We created one pool (of two samples) for assessment of parallelism and a second pool (adding seven additional samples) to assess the spike recovery. Sample information is given in the appendix (table A4).

### 2.3 Dog and human samples

Dog (*Canis lupus familiaris*) and human (*Homo sapiens*) urine pools were created through volunteer collections among laboratory staff/students and their pet dogs, as well as using extra volume collected for ongoing studies, under University of Arizona IACUC Protocol #16-175 and IRB Protocol #1808883345.

### 2.4 Solid-phase extractions

#### 2.4.1 Hydrophilic-Lipophilic Balanced (HLB)

Hydrophilic-Lipophilic Balanced reversed-phase extractions were conducted using the Oasis PRiME HLB 1 cc cartridges (Waters Corporation, Milford, MA, USA, Part Number: 186008055) and a positive pressure manifold (Biotage PRESSURE+48). All samples were diluted into an equal volume of 0.1% trifluoroacetic acid (TFA) in water, vortexed for at least 30 s, and then centrifuged. Exact speeds and times varied, in part due to sample volumes; all samples were centrifuged at either 10,000 rpm (RCF = 9632 x g) for 5 min or 5,000 rpm (RCF = 4,696 x g) for 15 min.

SPE cartridges were conditioned first with 1 ml acetonitrile (ACN) with 0.1% TFA, and then with 1 ml 0.1% TFA in water. The entire sample supernatant was loaded onto the cartridge. Each cartridge was then washed with 1 ml 10% ACN, 0.1% TFA. Finally, samples were eluted with 2 ml 30% ACN, 0.1% TFA. This procedure is identical to that reported in Gnanadesikan et al. (2021), although 2 ml of eluant was used in all cases, to maximize recovery.

#### 2.4.2 Mixed-mode Cation Exchange (MCX)

Mixed-mode Cation Exchange extractions were performed using Oasis PRiME MCX 1 cc cartridges (Waters Corporation, Milford, MA, USA, Part Number: 186008917) and a positive pressure manifold (Biotage PRESSURE+48). All samples were diluted into an equal volume of loading buffer (200 mM ammonium formate with phosphoric acid to achieve pH 5). Samples were vortexed for at least 30 s and then centrifuged, as above, at either 10,000 rpm (9632 x g) for 5 min or 5,000 rpm (4,696 x g) for 15 min.

The MCX cartridges were first conditioned with 1 ml methanol (MeOH) and 1 ml of deionized water (Thermo Scientific #751628). The entire supernatant of the sample was then loaded onto each cartridge. Cartridges were washed with successive 1 ml washes: 1 ml deionized water, 4 ml wash buffer (60% loading buffer, 40% MeOH), 1 ml deionized water. Finally, each sample was eluted in 2 ml of elution buffer (50% MeOH, 50% ammonium hydroxide solution). The pH of the elution buffer was 12 in the three-way kit comparison; the subsequently optimized protocol— used for the individual-level analyses—uses pH 11. We experimented with various combinations of loading, wash, and elution buffers and volumes to optimize this protocol, as reported in the appendix.

#### 2.4.3 All extracted samples

The eluted samples were centrifuged at 3,000 rpm (1690 *×* g) for 5 minutes to ensure that liquid on the side of the tubes settled at the bottom. All samples were stored at -80 °C overnight and lyophilized the next day using a CentriVap (Labconco model #7810016) with the following settings: CentriVap unheated, cold trap -80–85 °C, vacuum 0.3–0.4 mbar. Samples were reconstituted in assay buffer immediately before assay. Individual samples were reconstituted in 250 µl assay buffer; certain pools for multi-kit comparisons or validations used larger volumes, with concentration factors specified below. Reported concentrations are corrected for the concentration or dilution factors inherent in the extraction.

#### 2.4.4 Extraction efficiency

Extraction efficiencies were determined by extracting spiked buffer. We used the stock oxytocin solution from the Arbor Assays kit and diluted it to 1,000 pg/ml in assay buffer. Extractions were performed on 250 µl aliquots, while an additional aliquot was frozen for assay unextracted. Reported extraction efficiencies were calculated as (extracted measurement / unextracted measurement) *×* 100.

### 2.5 Oxytocin assays

Except where otherwise noted, oxytocin was measured using the Arbor Assays Oxytocin EIA kit (Catalog #K048-H5), following the recommended assay protocol (although not using the extraction solution provided). The reported sensitivity for this kit is 17.0 pg/ml and the lower limit of detection is 22.9 pg/ml; the limit of detection of a given experiment is dependent on (multiplied or divided by) the dilution or concentration factor inherent in the extraction, specified below. Arbor Assays reports that cross-reactivity is 94.3% for isotocin, 88.4% for mesotocin, and less than 0.15% for vasotocin and arginine vasopressin.

In the three-way kit comparison, we also used the Enzo Life Sciences (Catalog #ADI-900–153A-0001) and Cayman Chemical (Catalog #500440) kits. The manufacturer-reported sensitivities are 15 pg/ml for Enzo and 20 pg/ml for Cayman. The reported cross-reactivities for the Enzo kit are: mesotocin 7.0%, Arg^8^-vasotocin 7.5%, and *<* 0.02% for all other reported compounds including Arg^8^-vasopressin. Reported cross-reactivities for the Cayman kit are 100% for mesotocin and isotocin and *<* 0.01% for all other reported compounds including Arg^8^-vasopressin.

#### 2.5.1 Coefficients of variation

For unextracted samples of both genotypes (experiment 1), the mean intra-assay coefficient of variation (CV) between duplicates was 7.1%. For MCX-extracted samples (experiment 3), the mean intra-assay CV for samples that measured above the assay’s limit of detection (n = 6) was 7.4%. Results were not compared across plates (except for the multi-kit comparison).

### 2.6 Epitope mapping

Epitope mapping of the Arbor Assays antibody was conducted by Pepscan Presto BV (Lelystad, The Netherlands). The basic approach involves synthesizing peptide fragments and testing an antibody’s affinity for each sequence (Geysen et al., 1984). First, Pepscan constructed arrays of peptide fragments to assess the linear and conformational epitope binding; these arrays consisted of all possible 10-amino-acid fragments of oxytocin/neurophysin I prepropeptide, the larger precursor protein from which oxytocin is derived. Next, they performed fine-mapping of the epitope region by using single amino acid substitutions for each of the 9 amino acids in oxytocin to determine how each substitution affected antibody binding. The binding of antibody to each of the synthesized peptides was tested in a Pepscan-based immunoassay. This approach enables a comparison of the antibody’s affinity for different peptide sequences, which can shed light on the antibody’s specificity. Specifically, sequences that do not demonstrate high levels of antibody binding are unlikely to interfere with immunoassay measurement, while sequences that show binding levels similar to the identified epitope may cause interference.

### 2.7 Oxytocin fragments

Given that fragments of oxytocin are known to be formed through multiple metabolic processes (Uvnäs Moberg et al., 2019), we hypothesized that these fragments might cross-react with immunoassay antibodies. We focused on three biologically active fragments of oxytocin that have been studied previously: OT 2-9, OT 4-9, and OT 7-9, where the numbers indicate the residues of oxytocin (a nonapeptide) contained in the fragment. OT 7-9 is more commonly known as melanocyte-inhibiting factor 1 (MIF-1). In order to assess the cross-reactivities of these fragments on the Arbor Assays kit, we purchased MIF-1 from Cayman Chemical (Catalog #24476) and had OT 2-9 and OT 4-9 synthesized by Biomatik (Ontario, Canada). Since oxytocin is cyclic, with residues 1 and 6 connected by a disulfide bridge, OT 2-9 and OT 4-9 were synthesized with a cystine at position 6 (cysteine-cysteine, connected by a disulfide bridge), which is consistent with known metabolic pathways (Uvnäs Moberg et al., 2019). For more information on these fragments, see figure 4.

We reconstituted a single 10 mg aliquot of MIF-1 in 10 ml of deionized water, freezing reconstituted aliquots for future assays. The synthesized fragments came in 1 mg aliquots, so we used a new aliquot for each assay; we reconstituted each aliquot in 1 ml deionized water to create a 1 mg/ml solution, thus variation in these aliquots could contribute to variation in measured cross-reactivities across assays. On the day of each assay, the 1 mg/ml solution was diluted in assay buffer to the starting concentration of the assay. In a first experiment, we created 1,000,000 pg/ml starting solutions and then conducted five 1:10 part-to-whole (p:w) serial dilutions for each fragment to identify what orders of magnitude would result in measurements within the standard curve. Informed by these results, we conducted a second assay with serial dilutions targeting the middle of the standard curve. Reported mean cross-reactivities were calculated based on those dilutions that measured within 30-70% of the assay’s maximum binding (B_0_) as the percentage of the fragment’s known concentration that was measured by the assay. Although immunoassay results are usually reported by weight (e.g. pg/ml), since the fragments are very different sizes, we report cross-reactivities by both molarity and weight. The cross-reactivities-by-molarity represent a direct comparison based on the number of peptides present, while cross-reactivities-by-weight may be helpful in considering what fraction of a measurement might be due to each of these fragments.

In a third assay, we explored whether the HLB and MCX extractions removed these fragments. Each fragment was prepared to a concentration that we expected would result in approximately 30% of the maximum binding, based on the previous assays. An aliquot was frozen overnight to measure unextracted, and two aliquots were extracted using the HLB and MCX solid phase extraction methods. The extractions were performed on 250 µl of sample, the eluate was lyophilized, and the sample was reconstituted in 250 µl of assay buffer at the time of assay.

### 2.8 Analytical validations

Analytical validations were conducted with wildtype mouse, sifaka, dog, and human sample pools. Each species differed in the amount of sample extracted (and thus starting concentration/dilution factor) but were otherwise treated identically. Parallelism was assessed via serial dilution (2:3, p:w) of urine pools and calculation of the coefficient of variation on corrected concentrations at each dilution (Andreasson et al., 2015). Starting dilution/concentration factors for each parallelism were 1.33x for mouse and sifaka, 5.3x for dog, and 16x for human.

Spike recovery was assessed using samples at a single dilution (ratio of volume applied to the cartridge for SPE to the volume reconstituted): 2:5 for mouse, 1:2 for sifaka, 4x for dogs, and 8x for humans. All spike recovery extractions were reconstituted in 250 µl assay buffer; sample volumes were 100 µl for mouse, 125 µl for sifaka, 1 ml for dog, and 2 ml for humans. Spiked samples consisted of 90% sample matrix at the specified dilution and 10% synthetic oxytocin in assay buffer (kit standards 1–5). Percent recovery was calculated as (observed / expected) X 100, where the expected values were measured independently by adding each spike to assay buffer.

### 2.9 Statistical software

Statistical analyses were conducted in R version 4.1.1 (R Core Team, 2021). For the unextracted urine comparison in experiment 1, distributions were not significantly different from a normal distribution (Shapiro-Wilk’s W_*W T*_ = 0.948, *p*_*W T*_ = 0.644; *W*_*KO*_ = 0.955, *p*_*KO*_ = 0.677), so Welch’s t-test was used to compare the means. For the HLB extractions in experiment 2 and the three-way kit comparison in experiment 3, single measurements were made on pooled samples and no statistical tests were used. For the individual-level results in experiment 3, sample sizes were small (n = 6) for each genotype, so we used a Mann-Whitney test to avoid normality assumptions. See the appendix for a discussion of power analyses.

## 3 Results

### 3.1 Experiment 1: unextracted urine

Given our past success assaying unextracted mouse plasma samples (Gnanadesikan et al., 2021), we began by assessing the genotype contrast in unextracted diluted mouse urine samples on the Arbor Assays Enzyme Immunoassay (EIA) kit. Due to limited sample volume for some individuals, some samples were assayed at a 1:10 (p:w) dilution into Arbor Assay Buffer, while others were measured at a 1:25 dilution. Based on a serial dilution of a single wildtype C57BL/6J urine sample, we expected both of these dilutions to be within range of the assay (see appendix, table A1).

Knockout samples were indistinguishable from wildtype samples (Mean ± SD: *WT* = 930.8 ± 512.7 pg/ml, KO = 885.7 ± 295.6 pg/ml; Welch’s *t*(13.5) = 0.248, *p* = 0.808; figure 1). We interpreted this as indicating a high level of interference in the immunoassay and proceeded to explore various extraction options to eliminate or minimize this interference.

**Figure 1:**
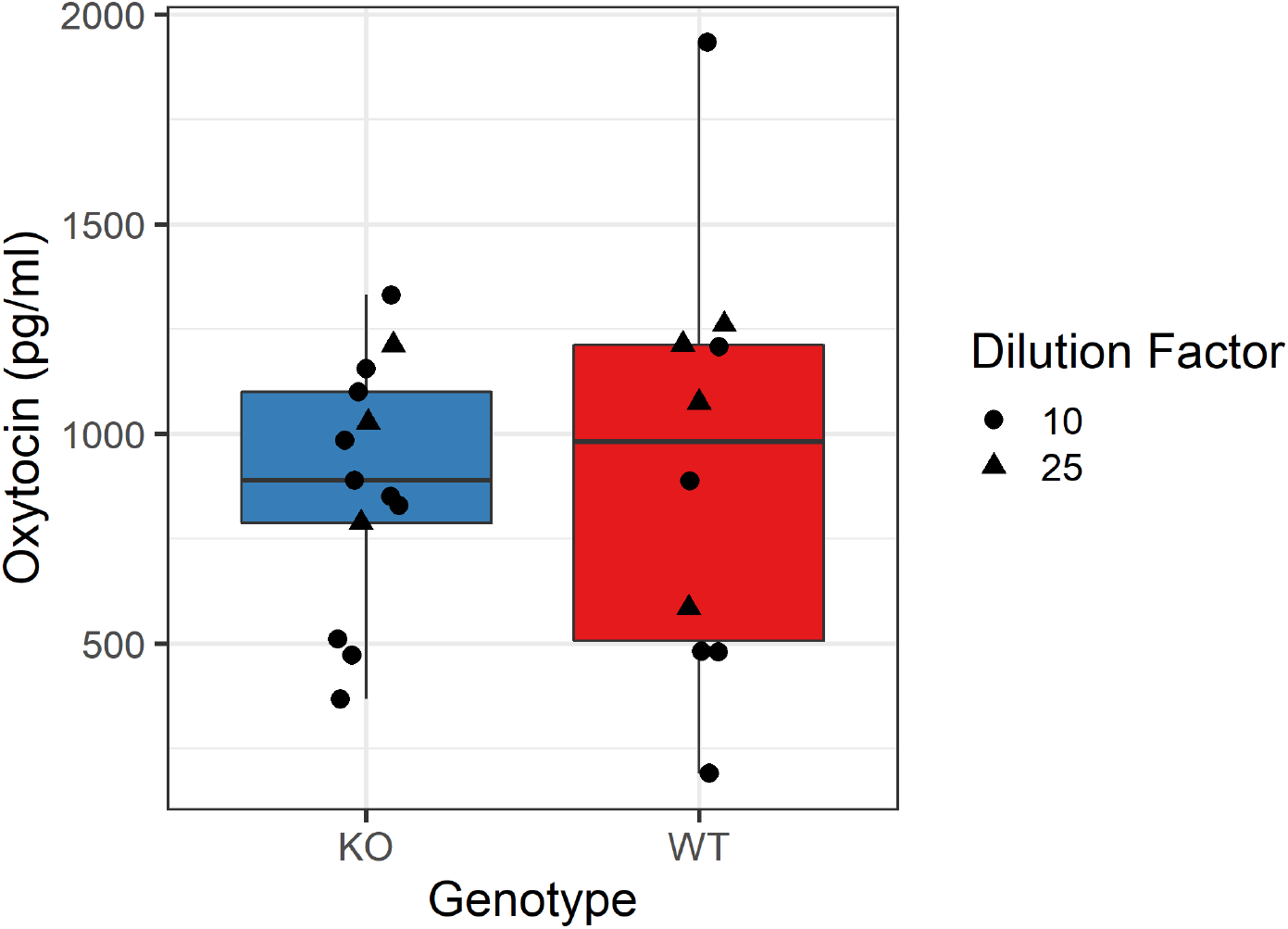
Boxplot comparing the dilution-corrected measured oxytocin concentrations in unextracted urine samples across genotypes. Depending on available sample volume, samples were assayed at a 1:10 (circle) or 1:25 (triangle) dilution into assay buffer. The mean measured oxytocin concentrations were not significantly different across genotypes (Mean ± SD: WT = 930.8 ± 512.7 pg/ml, KO = 885.7 ± 295.6 pg/ml; Welch’s *t*(13.5) = 0.248, *p* = 0.808). Note that this pattern was true even with the presence of a single high WT sample; excluding this point, Mean WT = 819.6 ± 395.5, which is below the KO mean. The limits of detection are 229 pg/ml for samples assayed at a 1:10 dilution and 573 pg/ml for those assayed at a 1:25 dilution.

### 3.2 Experiment 2: basic HLB extraction

Since solid-phase extraction is commonly used to remove interfering molecules from biological matrices, we evaluated the performance of a relatively simple hydrophilic-lipophilic balanced (HLB) extraction. We had previously validated this method for mouse plasma (Gnanadesikan et al., 2021). With urine samples, this extraction resulted in a clear genotype contrast, but there was still considerable interference in the knockout samples (WT_*pool*_ = 611.9 pg/ml; KO_*pool*_ = 188.03 pg/ml). Subsequent replications found similar results (not reported). We assessed whether an increased wash volume would eliminate the observed interference, but instead found that we lost signal in both genotype pools, and the genotype contrast was not improved (WT_*pool*_ = 116.49 pg/ml; KO_*pool*_ = 72.81 pg/ml).

### 3.3 Experiment 3: improved MCX extraction method

In order to improve our extraction selectivity, we developed and optimized a new method using a mixed-mode cation exchange (MCX) cartridge. This enabled us to manipulate the retention using both pH and organic content of the loading, wash, and elution buffers. For details of the methods development, see the appendix (section 3, figures A1–A5).

Using this new method, we then compared the MCX and HLB extraction methods across three different oxytocin immunoassay kits, hypothesizing that differences in the kit chemistries— especially antibodies—might lead to different susceptibilities to interference. We found that the HLB extraction was susceptible to varying levels of interference across kits, but the MCX extraction minimized this interference on all three kits (KO measurements all *<* 40 pg/ml, corrected concentrations *<* 70 pg/ml; WT measurements 124-171 pg/ml, corrected concentrations 206-286 pg/ml; figure 2).

**Figure 2:**
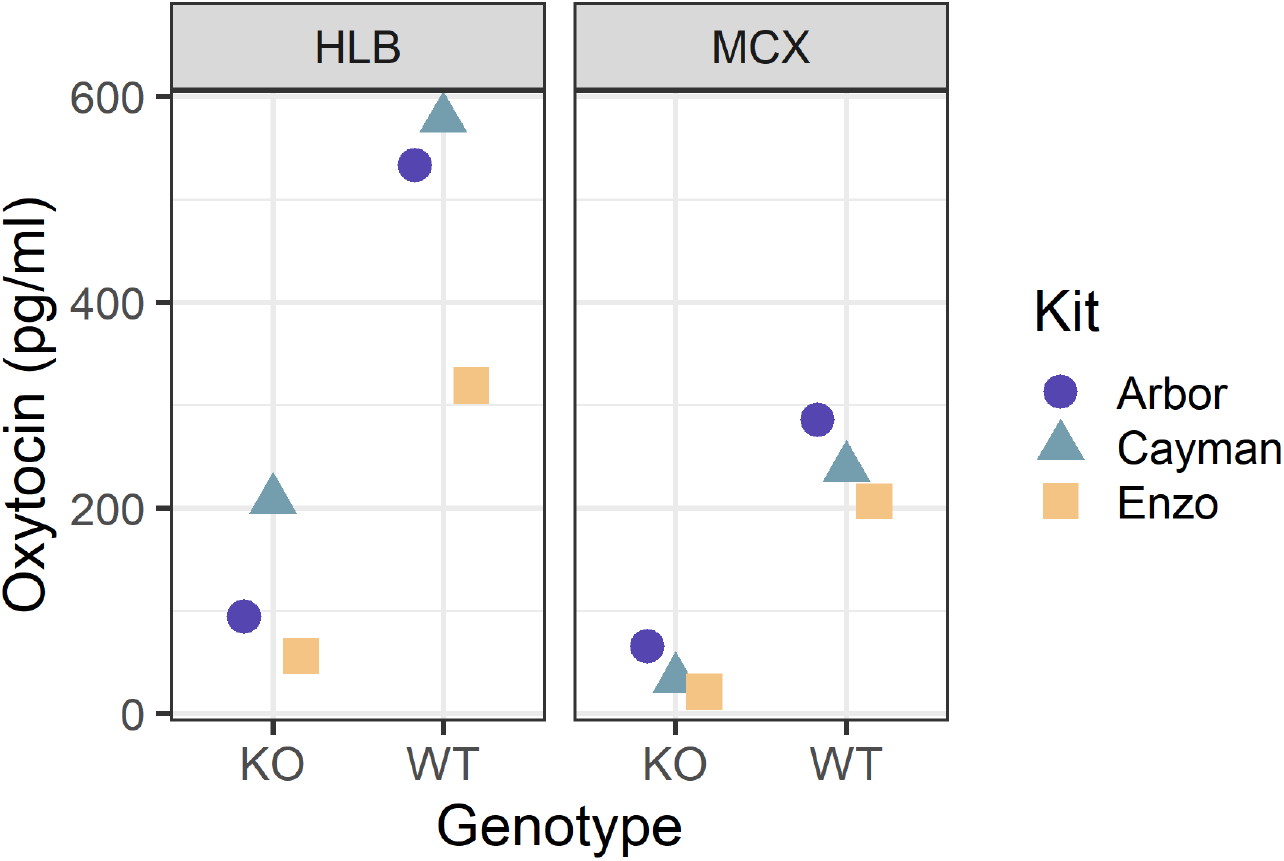
One pooled sample for each genotype was measured on three commercial kits after either an HLB or MCX solid phase extraction. HLB samples demonstrated varying levels of interference on all kits. The MCX extraction minimized this interference and resulted in relatively concordant results across kits.

While these results were much improved, we wanted to further minimize the KO interference, with the aim of increasing the WT:KO ratio. We achieved this on the MCX by decreasing the pH of the elution buffer from 12.0 to 11.0. We then tested individual samples (some pooled) to examine the genotype contrast. All knockout samples measured below the assay’s limit of detection. As such, these values should not be interpreted (and in some cases had high CVs due to low measurement); nevertheless, the difference between groups is statistically significant (Mann-Whitney W = 0, p = 0.002; figure 3).

**Figure 3:**
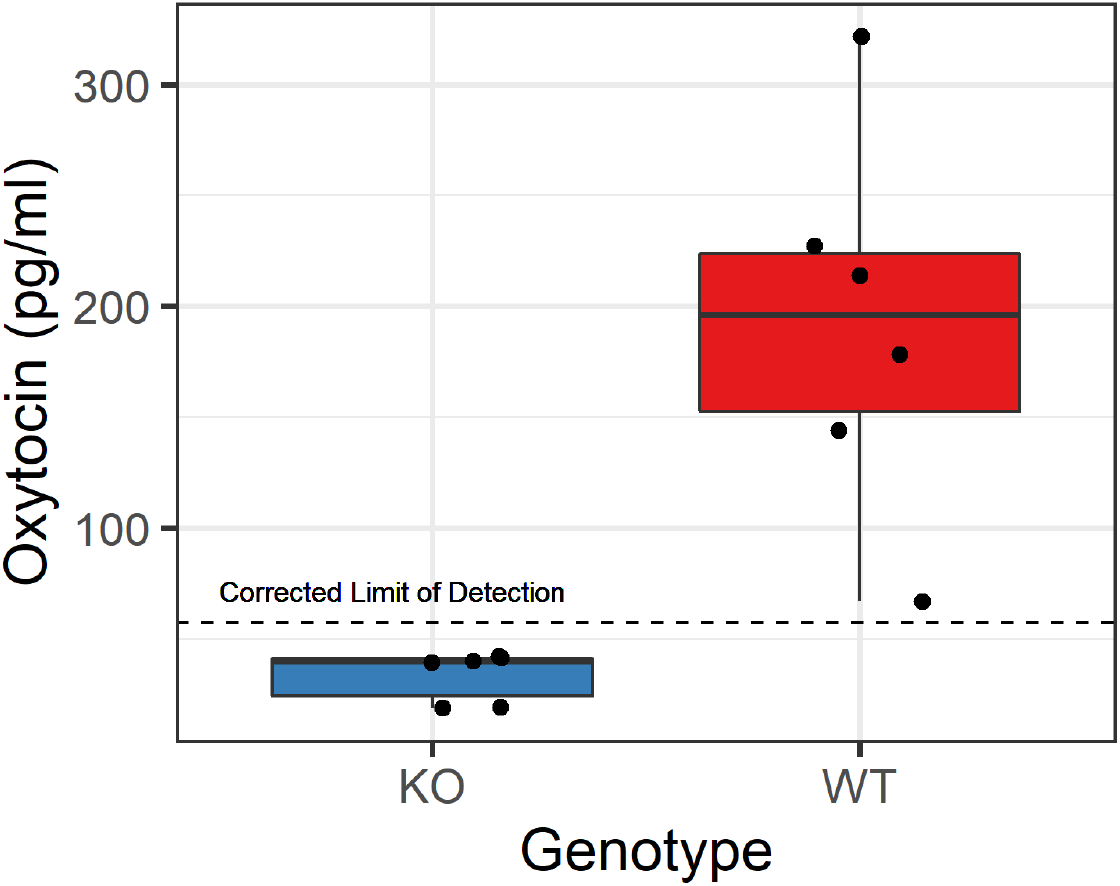
Genotype contrast using a more selective mixed-mode cation exchange SPE chemistry. Each point represents an individual mouse sample, some of which were pooled across 2-3 samples within the same (blinded) genotype code, due to limited sample volume, resulting in six samples per genotype. A clear genotype distinction can be seen between knockout (KO) and wildtype (WT) samples (Mann-Whitney W = 0, p = 0.002).

Using spiked buffer samples, we also assessed the extraction efficiency of the MCX method; in one assay across three samples, the extraction efficiency was 83%, while in another assay across two samples, the extraction efficiency was 92%.

### 3.4 Epitope mapping

Given the considerable interference that we observed in unextracted and HLB-extracted urinary samples, we decided to further probe the properties of the anti-oxytocin polyclonal rabbit antibody used in the Arbor Assays EIA. We therefore worked with Arbor Assays and Pepscan to perform both linear and conformational epitope mapping of this antibody. Epitope mapping tests an antibody’s affinity to a library of synthesized peptide fragments, thereby determining its epitope(s) and assessing its specificity.

The epitope mapping results identified a single epitope within oxytocin/neurophysin I prepropeptide, corresponding to residues 2-9 of oxytocin (YIQNCPLG). Further testing of this region— by assessing single amino acid substitutions—revealed that the core epitope is PLG (figure 4, A6). Almost any change to these residues results in highly disrupted binding, although the magnitude of this change was dependent on the exact substitution; substitutions outside of this core region had less impact on antibody binding (see supplementary data file for raw epitope mapping results).

**Figure 4:**
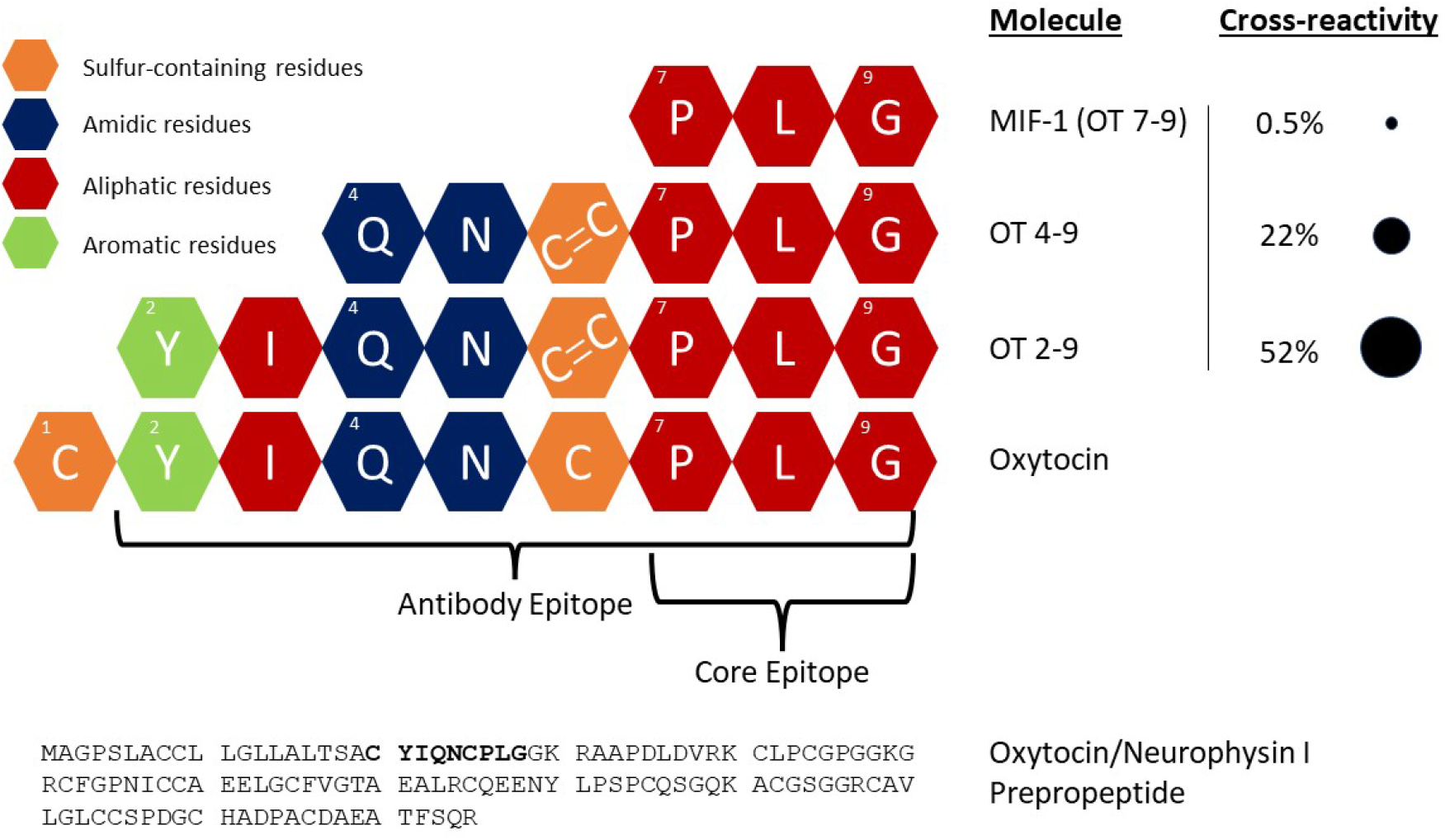
Oxytocin is a nonapeptide formed from the cleavage of oxytocin/neurophysin I pre-propeptide. Our epitope mapping of the Arbor Assays anti-oxytocin antibody demonstrates that the epitope spans oxytocin residues 2-9, while the core epitope consists of residues 7-9. We purchased three biologically active fragments of oxytocin to test their cross-reactivity with the Arbor Assays antibody, finding high cross-reactivity for 2-9, intermediate cross-reactivity for 4-9, and low cross-reactivity for 7-9 (also known as MIF-1). Cross-reactivities reported here are by weight (see text for cross-reactivities by molarity). Since oxytocin is cyclic, with residues 1 and 6 connected by a disulfide bridge, OT 2-9 and OT 4-9 were synthesized with a cystine at position 6 (cysteine-cysteine), consistent with known metabolic pathways.

The linear and loop arrays produced similar results, as did versions allowing the free cysteines to bridge, suggesting that the conformational structure of oxytocin—including the disulfide bridge between the cysteine residues—is not important for epitope recognition, consistent with earlier studies reporting immunoreactivity with the linear form of oxytocin after a reduction/alkylation procedure (Brandtzaeg et al., 2016).

### 3.5 Oxytocin fragment cross-reactivities and extractions

Given what we learned about the epitope of the Arbor Assays antibody, and especially that the core epitope consists of residues 7-9, we investigated the cross-reactivity of this antibody with known, biologically active fragments of oxytocin (Uvnäs Moberg et al., 2019). Specifically, we purchased OT 2-9, OT 4-9, and OT 7-9, the latter more commonly known as melanocyte-inhibiting factor 1 (MIF-1). Across multiple dilutions, the average cross-reactivities-by-molarity, measured within the range of 30-70% of the assay’s maximum binding (B_0_) were 52% for OT 2-9 (also 52% by-weight), 22% for OT 4-9 (32% by weight), and 0.5% for MIF-1 (2% by weight; figures 4 and 5; table A5).

**Figure 5:**
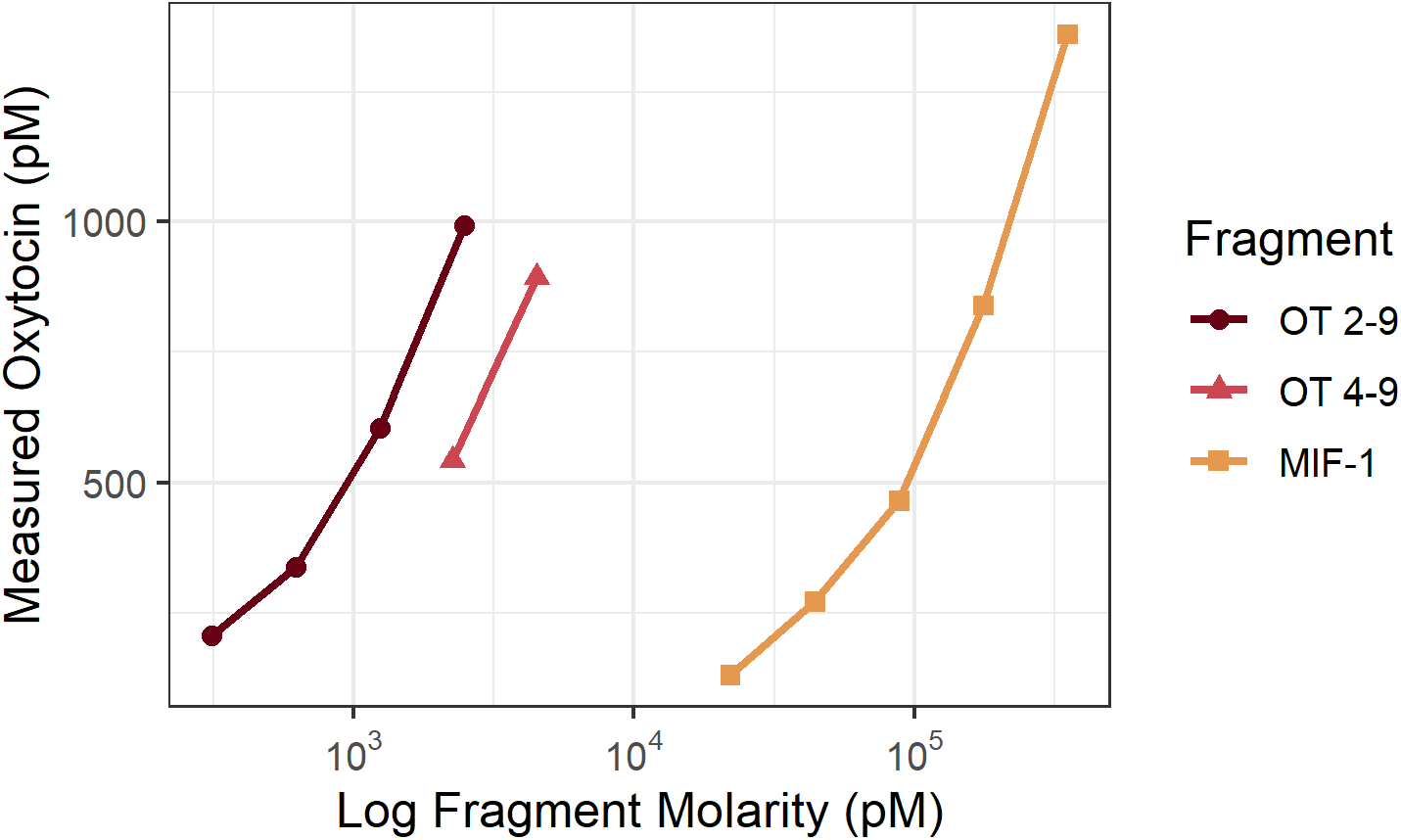
Measured oxytocin concentrations on the Arbor Assays EIA kit vs. known concentrations of three biologically active fragments of oxytocin (log scale). Only measurements within the range of 30-70% of the assay’s maximum binding are shown and used to calculate cross-reactivity, the percent of the known fragment concentration that is measured as oxytocin. OT 2-9, the fragment most similar to oxytocin, had the highest cross-reactivity (52% by-molarity and by-weight), followed by OT 4-9 (22% by-molarity, 32% by-weight). MIF-1, the smallest fragment tested, displayed the lowest cross-reactivity (0.5% by-molarity, 2% by-weight).

Finally, we tested both the HLB and refined MCX extractions on each of the three oxytocin fragments, as well as oxytocin itself. While both extractions retained ¿ 90% of oxytocin itself, extraction considerably reduced the measured oxytocin concentrations for all three fragments. Both extractions removed most of the fragments, reducing the assay’s measurement by 1-2 orders of magnitude. The extractions were more effective at eliminating OT 4-9 and MIF-1; for OT 2-9—the molecule most similar to oxytocin—the MCX extraction removed considerably more of the fragment (measured 117.15 pg/ml) than did the HLB extraction (measured 283.05 pg/ml; see figure 6).

**Figure 6:**
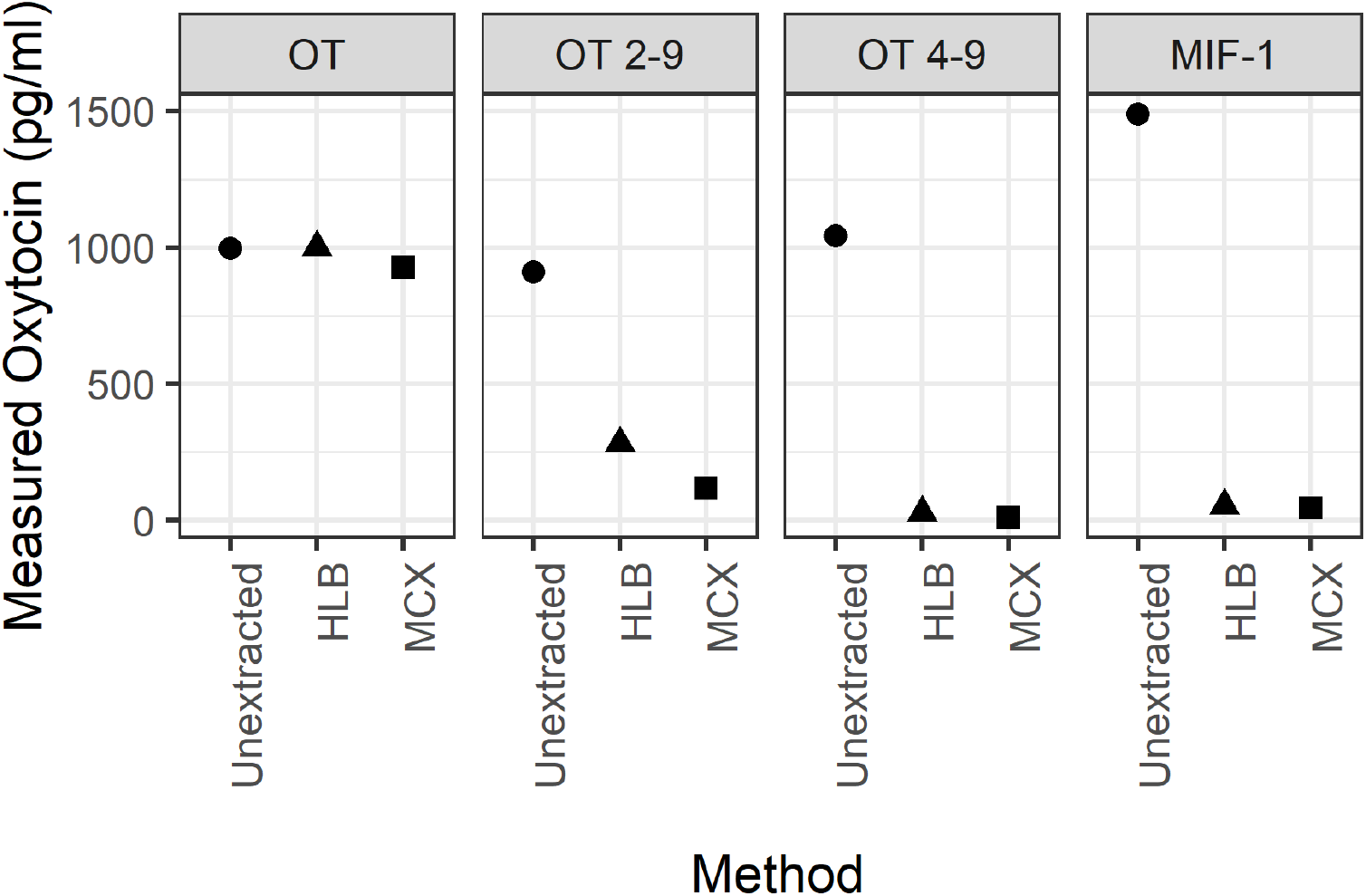
Oxytocin and each of the three oxytocin fragments was measured with no extraction and after one of two solid-phase extractions: HLB or MCX. Both extractions retain most of the oxytocin present in spiked buffer (HLB: 100%, MCX 93%). However, the extractions eliminate most of the signal measured for each fragment of oxytocin, with the MCX extraction removing more of oxytocin 2-9 than the HLB extraction. Note that the actual starting concentrations for each fragment are higher than their measured concentrations (OT 2-9: 2,500 pg/ml; OT 4-9: 4,000 pg/ml; MIF-1: 100,000 pg/ml).

### 3.6 Analytical validations: mouse, sifaka, dog, and human

Using the optimized MCX method, we conducted analytical validations on pooled samples from four species/strains: wildtype mice (*Mus musculus*), Verreaux’s sifaka (*Propithecus verreauxi*), domestic dogs (*Canis lupus familiaris*), and humans (*Homo sapiens*).

The wildtype mouse pool diluted parallel to the standard curve (CV of corrected concentrations = 10.1%; figure A8) and exhibited excellent spike recovery for samples measured at a 2:5 dilution factor (mean ± SD = 114 ± 7%). Similarly, pooled sifaka urine samples demonstrated good parallelism (CV of corrected concentrations = 17.5% over full range; 5.11% for lower 5 dilutions; figure A9) and excellent spike recovery measured at a 1:2 dilution (mean ± SD = 114 ± 6%). A dog urine pool diluted extremely parallel to the standard curve (CV of corrected concentrations = 6.0%; figure A10) and exhibited good spike recovery when assessed at a 4x concentration (mean ± SD = 119 ± 9%).

Lastly, a human urine pool diluted relatively parallel to the standard curve, especially for higher concentrations (figure A11). The CV of corrected concentrations over the entire range (16x-2.1x) was 18%; excluding the fourth dilution, which had a duplicate CV of 25%, this rose to 19%. However, the last two dilutions measured somewhat higher than expected, thus for the first three dilutions alone, the CV of corrected concentrations was 6.6%. Human urine also exhibited good spike recovery when measured at an 8x concentration (mean ± SD = 123 ± 8%). It should be noted, however, that the specific gravity for this sample pool is 1.023 and had a corrected concentration of approximately 7 pg/ml. Higher concentration factors may be necessary for more dilute samples. More experimentation is needed to determine whether an ideal protocol exists for samples of all concentrations, or what the best methods are for compensating for the wide variation in specific gravity (and consequently oxytocin concentrations) in human urine. Relatedly, in some subsequent experiments, spike recovery for certain human urine pools was above acceptable thresholds, so further optimization may be necessary for human samples.

## 4 Discussion

Our results suggest that many existing urinary oxytocin methods may be suboptimal. We found that unextracted urine samples—measured at a 1:10 or 1:25 dilution in assay buffer—were indistinguishable on the Arbor Assays kit across oxytocin knockout and wildtype mice. A basic reversed-phase SPE on the Waters Oasis PRiME HLB cartridge improved the genotype contrast, but still exhibited considerable interference, which was observed at varying levels across three different commercially available immunoassay kits. This interference was eventually minimized with the development of a new SPE method using the Waters Oasis PRiME MCX cartridge, which utilizes a mixed-mode cation exchange chemistry. Although we have not tested all of the cartridge-extraction-kit combinations that have previously been used in the literature, our findings across three different commercially available kits suggest that there is strong potential for interference in immunoassays of urinary oxytocin, even following sample extraction. It is notable that the concentrations we report are lower than those reported with some other methods. For example, Wirobski, Range, et al. (2021) reported pet dog concentrations in the range of 77 – 1500 pg/ml (Mean ± SD = 440 ± 290) before correcting for specific gravity (SG), and 130 – 950 pg/ml (Mean ± SD = 430 ± 190) after doing so. The dog pool that we used for both parallelism and spike recovery analyses had concentrations of 40 – 50 pg/ml (no SG correction), and in another study in progress, pet dog samples have generally measured *<* 100 pg/ml before SG-correction, and typically *<* 250 pg/ml after SG-correction.

In methods development work, higher concentrations are often assumed to be better (e.g. higher extraction efficiency or improved storage). This assumption is tempting, in part because many studies suffer from an inability to detect analytes at low concentrations. However, this assumption may lead us astray without a careful consideration of potential sources of interference, which may be particularly abundant in urine. On the other hand, it has also been assumed—without sufficient empirical evidence—that high oxytocin concentrations are biologically implausible. This and other recent work (Gnanadesikan et al., 2021) suggest that some methods, at least, are not susceptible to interference in mouse samples.

Feldman et al. (2011) found that in human samples, while plasma and salivary measures of oxytocin were correlated, extracted urinary measurements were not. Weber et al. (2017) also found no correlation between unextracted plasma and unextracted urinary oxytocin measurements in premature human infants. While there are many possible reasons for these findings, including that different biological matrices are thought to integrate over different periods of time, another plausible interpretation in light of our results is that the urine measurements in these studies were susceptible to interference. This interpretation is supported by studies that have found correlations between plasma and extracted urinary oxytocin in certain conditions and time windows (Amico et al., 1987; Francis et al., 2016).

It should be noted that many of the previously demonstrated associations between urinary oxytocin measurements and social behavior may be robust, despite these potential limitations, especially if employing a change-from-baseline approach. Nevertheless, many studies may have inflated concentrations due to the type of interference we demonstrated here. This interference may also contribute to negative findings if the noise introduced by interference obscures an underlying effect, which may be especially important for studies of individual differences.

While the success of our method across a wide variety of mammalian species—including members of the orders Primates, Rodentia, and Carnivora—is promising, the extraction method as reported here may not be optimal for all species. It should be noted, however, that this method worked well even in wild sifaka samples, which are highly concentrated due to their extremely limited water intake; therefore we do not expect highly concentrated urine to be problematic. Nevertheless, we strongly encourage anyone who aims to extend this work to other species and populations to perform a careful analytical validation with their own samples, particularly parallelism and spike recovery, although these tests alone may not be sufficient to determine the absence of interference.

Our epitope mapping results illuminate one of the core components of any immunoassay: the antibody. The importance of the core epitope—spanning residues 7-9—is consistent with the manufacturer’s reports of low cross-reactivity with mesotocin and vasopressin, since both of these related nonapeptides differ from oxytocin at the eighth residue. It is worth noting that a novel form of oxytocin has been documented in some species of New World monkeys; this form of oxytocin has a substitution at residue 8, with proline replacing the ancestral leucine (Lee et al., 2011). Given our epitope mapping results, we would expect binding for this form of oxytocin to be significantly reduced. Interestingly, although the identity of each residue in the rest of oxytocin (1-6) is less important for antibody binding, cross-reactivity drops rapidly for smaller fragments.

This demonstrated cross-reactivity is important for several reasons. First, measurements of oxytocin by immunoassay likely include some amount of signal from these bioactive fragments; this interpretation is supported by the observation that injection of radiolabelled oxytocin (in marmosets) results in urinary samples that produce multiple radioactive fractions after HPLC, including ones other than the standard (Seltzer & Ziegler, 2007). Second, although the cross-reactivity for MIF-1 seems low, it is a very small fragment, with a sequence that is common in a variety of peptides. This cross-reactivity may partially explain why immunoassay of urinary samples is more difficult than for plasma samples, given that urine is a waste product, full of thousands of known and unknown metabolites (Bouatra et al., 2013). A BLASTP (Altschul et al., 1990) search of the NCBI Protein Reference Sequences database (O’Leary et al., 2016), restricted to *Homo sapiens*, on oxytocin 4-9 reveals 678 unique sequences with 100% identity (i.e. containing QNCPLG), and many more with partial identity. This result suggests that a large number of metabolites might have sufficient sequence similarity to cross-react with the Arbor Assays anti-oxytocin antibody, although we do not know which specific fragments are present in urine or what metabolic processes produce them. Importantly, oxytocin has been shown to be one of the least abundant—while still detectable—metabolites in human urine, while other metabolites can be more abundant by up to 10 orders of magnitude (Bouatra et al., 2013). Thus, even peptide fragments with low cross-reactivities could interfere when present in high enough concentrations.

It should also be noted that while other immunoassay kits use different antibodies, given that the oxytocin molecule is so small, other antibodies likely have a similar susceptibility to interference, even if their epitope and/or core epitope is slightly different. For example, we previously demonstrated that the Enzo Life Sciences kit is susceptible to interference with plasma samples in situations where the Arbor Assays kit is not, presumably reflecting differences in their epitopes or specificity (Gnanadesikan et al., 2021). Similarly, Wirobski, Schaebs, et al. (2021) found that the Arbor Assays kit performed better than the Enzo kit for human and canine urinary oxytocin, with better spike recoveries, lower intra-assay CVs, and increased sensitivity. Nevertheless, given the fundamental constraint posed by the small size of the oxytocin molecule, improved immunoassay of urine samples will most likely rely on improved extraction techniques, such as the solid-phase extraction method that we report here.

Whether immunoreactivity from oxytocin’s metabolites should be considered interference depends on the perspective of the end user. Indeed, analytes measured in excretory products such as urine and feces often involve binding to metabolites of the target analyte rather than the parent molecule, and in some cases this may be desirable (Palme et al., 2013). It has recently been proposed that many of oxytocin’s functions may derive from active fragments of the nonapeptide (Uvnäs Moberg et al., 2019), suggesting that oxytocin metabolites may be relevant biomarkers for some aspects of the oxytocinergic system. Our results indicate that several bioactive fragments of oxytocin are nearly entirely eliminated using solid-phase extraction, making this an important consideration for the protocols described here.

Some oddities of urinary oxytocin measurement have been reported previously. Schaebs et al. (2019) performed a validation of dog and wolf samples on the Enzo kit, but found that despite good parallelism, the spike recoveries and extraction efficiencies were concerningly high. Reyes et al. (2014) reported that 18% of their samples produced out-of-range high measurements on the Enzo Life Sciences kit even after solid-phase extraction. Interestingly, when assayed at a two-fold dilution, not only were the samples within range, but the corrected oxytocin concentrations were also within range, suggesting some form of interference when assayed at higher concentrations. Reyes et al. (2014) hypothesize that this may be due to dehydration and overly concentrated samples, supported by the finding that out-of-range samples had higher creatinine values. Higher creatinine was also associated with higher vasopressin measurements, higher specific gravity, and more acidic samples. Our work suggests that either high concentrations of metabolites or the altered pH of dehydrated samples could explain this result, and the MCX method—with the pH of buffers optimized for both retaining oxytocin and minimizing interference—improves assay performance considerably.

Lastly, we would like to caution that this and our previous work (Gnanadesikan et al., 2021) emphasize the need for detailed reporting of immunoassay methods, not only in validation papers, but also accompanying any empirical result. This reporting should include the details of extraction methods and immunoassay kits, sample and buffer volumes, dilution factors, and other aspects of the sample treatment and processing protocols. Attention to these details will aid in both interpretation and replication of reported results.

## 5 Conclusions

Our immunoassay results indicate that many existing methods of measuring urinary oxytocin may be susceptible to interference. Our epitope mapping and fragment cross-reactivity results suggest this may be attributable to the abundance of metabolites in urine. Nevertheless, we show that a more selective extraction method, utilizing a mixed-cation exchange (MCX) chemistry, is successful at minimizing this interference. This method has the potential to improve urinary oxytocin studies in a wide variety of species and to open doors for reconsidering the relationships between urinary oxytocin and other physiological and behavioral measures.

## Supporting information

Epitope Mapping Data

## CRediT authorship contribution statement

**Gitanjali E. Gnanadesikan**: Conceptualization; Data curation; Formal analysis; Funding acquisition; Investigation; Methodology; Visualization; Writing – original draft. **Elizabeth A.D. Hammock**: Conceptualization; Resources (mouse samples); Writing – original draft. **Stacey R. Tecot**: Conceptualization; Funding acquisition; Resources (laboratory, sifaka samples); Project administration; Writing – original draft. **Rebecca J. Lewis**: Resources (laboratory, sifaka samples); Writing – review & editing. **Russ Hart**: Conceptualization; Funding acquisition; Methodology; Writing – review & editing. **C. Sue Carter**: Conceptualization; Writing – review & editing. **Evan L. MacLean**: Conceptualization; Funding acquisition; Methodology; Project administration; Resources (laboratory equipment); Writing – original draft.

## Acknowledgements

We would like to acknowledge the Pepscan team for their work on the epitope mapping, as well as Arbor Assays for their contributions. In particular, Bobbi O’Hara at Arbor Assays was extremely helpful at multiple stages of this work. We also benefited significantly from conversations with Terence Gorman at Waters, who advised us throughout the process of developing and optimizing the MCX method. Sifaka samples were collected and transported with funding by the National Science Foundation under award number BES-1719654, with the permission of CAFF/CORE, the Ministry of Water and Forests, and Madagascar National Parks, and the assistance of the University of Antananarivo, MICET, the Ankoatsifaka Research Station, the Sifaka Research Project staff, and Meredith Lutz, Diary Razafimandimby, Fanomezantsoa Razafimalala, Mc Antonin Andriamahaihavana, Sylvia Rahobilalaina, Hannah Carbonneau, and Allison Hays. This material is based upon work supported by the Eunice Kennedy Shriver National Institute of Child Health & Human Development of the National Institutes of Health under award number R21HD095217, by the National Institute of Mental Health under award number MH114994, and by the National Science Foundation Graduate Research Fellowship Program under grant number DGE-1746060. Any opinions, findings, and conclusions or recommendations expressed in this material are those of the authors and do not necessarily reflect the views of the National Institutes of Health or the National Science Foundation.

