## Appendix for "What are oxytocin assays measuring? Epitope mapping, metabolites, and comparisons of wildtype & knockout mouse urine"

### What are oxytocin assays measuring?
### Appendix: supplementary methods and results

Gitanjali E. Gnanadesikan, Elizabeth Hammock, Stacey R. Tecot, Rebecca J. Lewis, Russ Hart, C. Sue Carter, Evan L. MacLean

March 1, 2022

#### Contents

**Contents** 1

**1 Power analyses** 2

**2 Unextracted mouse urine dilutions** 2

**3 Methods Development** 3

**4 Epitope Mapping** 9

**5 Oxytocin Fragment Cross-Reactivities** 10

**6 Parallelism Plots** 11

**7 Data and Sample Information** 14

**8 References** 18

### 1 Power analyses

Informed by our previous results with plasma (Gnanadesikan et al., 2021), we ran a power analysis for a t-test using the *samplesize* package (Scherer, 2016) in R (R Core Team, 2021) and the following values: 90% power, alpha of 0.05, mean difference of 300 pg/ml, and standard deviations of 100 (WT) and 10 (KO). This produced a suggested sample size of 4 WT individuals and 2 KO individuals. Considering the possibility that concentrations in urine would be lower, we used the same power and alpha thresholds with a mean difference of 100 pg/ml, and standard deviations of 50 (WT) and 10 (KO). This produced a suggested sample size of 7 WT individuals and 2 KO individuals. Nevertheless, we pursued a more balanced design and began with a larger sample size for experiment 1 (WT: 10; KO: 14), especially given that we expected the knockout samples might be susceptible to interference. Experiment 3 used only six samples per genotype, which was in between these recommendations; however, since we were primarily interested in determining whether the knockout samples were below the assay's lower limit of detection, we proceeded with the samples we had available.

### 2 Unextracted mouse urine dilutions

To begin, we used a single wildtype (B6) sample and measured it at five dilutions to assess parallelism and dilutions for subsequent assays. The results did not quite meet standards for parallelism (CV of corrected concentrations = 24.5%), showing particularly high readings with more concentrated samples (i.e. dilutions 1 and 2). From the measured concentrations (see table A1), we determined that a 1:10 dilution for individual samples should measure within range, with a 1:25 dilution potentially working for samples with small volumes.

**Table A1:** A single wildtype B6 sample measured at 5 dilutions. Corrected concentrations were calculated by dividing by the noted dilution factor. The CV of corrected concentrations was 24.5%, and a decreasing trend can be seen in the corrected concentrations.

| **Dilution** | **Meas. OT (pg/ml)** | **CV (%)** | **Dilution Factor** | **Corr. OT (pg/ml)** |
|---|---|---|---|---|
| 1 | 814.507 | 1.15 | 0.36 | 2262.521 |
| 2 | 370.153 | 2.28 | 0.18 | 2056.404 |
| 3 | 131.066 | 2.04 | 0.09 | 1456.293 |
| 4 | 68.034 | 4.88 | 0.05 | 1511.872 |
| 5 | 28.996 | 4.74 | 0.02 | 1288.695 |

### 3 Methods Development

We primarily explored two solid-phase extraction (SPE) methods, using two different cartridges from Waters, the Hydrophilic-Lipophilic-Balanced (HLB) and Mixed Cation Exchange (MCX) cartridges. To develop each method, we first explored sequential elutions of synthesized oxytocin in buffer to determine under what conditions oxytocin elutes from each cartridge. This is quite straightforward on the HLB cartridges, because there is only one variable—organic content, here acetonitrile (ACN) specifically—to manipulate. For the MCX cartridges, this requires manipulating both organic content—here, methanol (MeOH)—and pH.

#### 3.1 HLB Extraction

For the HLB elution experiment, we loaded a single HLB cartridge with approximately 250 pg of oxytocin standard (at approximately 1,000 pg/ml concentration, measured unextracted as 1,028 pg/ml). We then collected the flow through sample, washed the cartridge with 1 ml 0.1% TFA in $H_2O$, and finally sequentially eluted with 1 ml of elution buffers of increasing ACN content (steps of 10%). We lyophilized/evaporated all samples in 250 µl of assay buffer and assayed them on the Arbor Assays kit. No measurable oxytocin eluted at 10%, the majority eluted at 20%, and some additional oxytocin eluted at 30% (see figure A1A). From this, we concluded that we could wash with $\leq$ 10% ACN without losing oxytocin (but potentially washing away some molecules that would not be washed away with water), and that eluting with 30% ACN would be sufficient to elute all the oxytocin in our samples (but without eluting too many other molecules that might interfere). This is the HLB method that was used for mouse plasma Gnanadesikan et al. (2021); further refinements of the wash step were later attempted for urine (see below).

#### 3.2 Phenomenex StrataX

We did also test the Phenomenex StrataX Polymeric Reversed Phase cartridge (Part #: 8B-S129-TAK) and found an elution profile similar to the HLB: no oxytocin eluted at 10% ACN, the entire expected oxytocin sample eluting at 20%, with a (smaller) residual elution at 30% ACN (see figure A1). We did not pursue further methods development with this cartridge, primarily due to the helpfulness of the Waters technical support team; however, it is worth noting that the elution profile was very similar to that on the Waters HLB cartridge, and recovery was perhaps even slightly higher.

#### 3.3 MCX Extraction

For the MCX cartridge, we started with the following procedure, after consulting with technical support at Waters: condition with 1 ml MeOH, wash with 1 ml $H_2O$, load the sample in an equal volume of loading buffer (200 mM $NH_4HCO_2$, 4% $H_3PO_4$), and wash with 1 ml wash buffer (30% MeOH, 70% loading buffer) followed by 1 ml $H_2O$. We then performed a sequential elution, with elution buffers that combined MeOH and a 5% aqueous solution of $NH_4OH$, in 10% steps from 40% to 90% MeOH. Most of the oxytocin was recovered in the first elution (40% MeOH), with some residual elution at 50% (see figure A1C). It is worth noting that the flow through and wash steps cannot be assayed, as the MCX loading buffer interferes with the immunoassay. Based on these results, we decided to elute at 50% MeOH.

Given that we had succeeded at retaining and eluting oxytocin, the next step was to refine the elution. Higher organic content and higher pH would both be expected to elute more molecules; therefore we aimed to identify the lowest pH for the elution buffer that would successfully elute

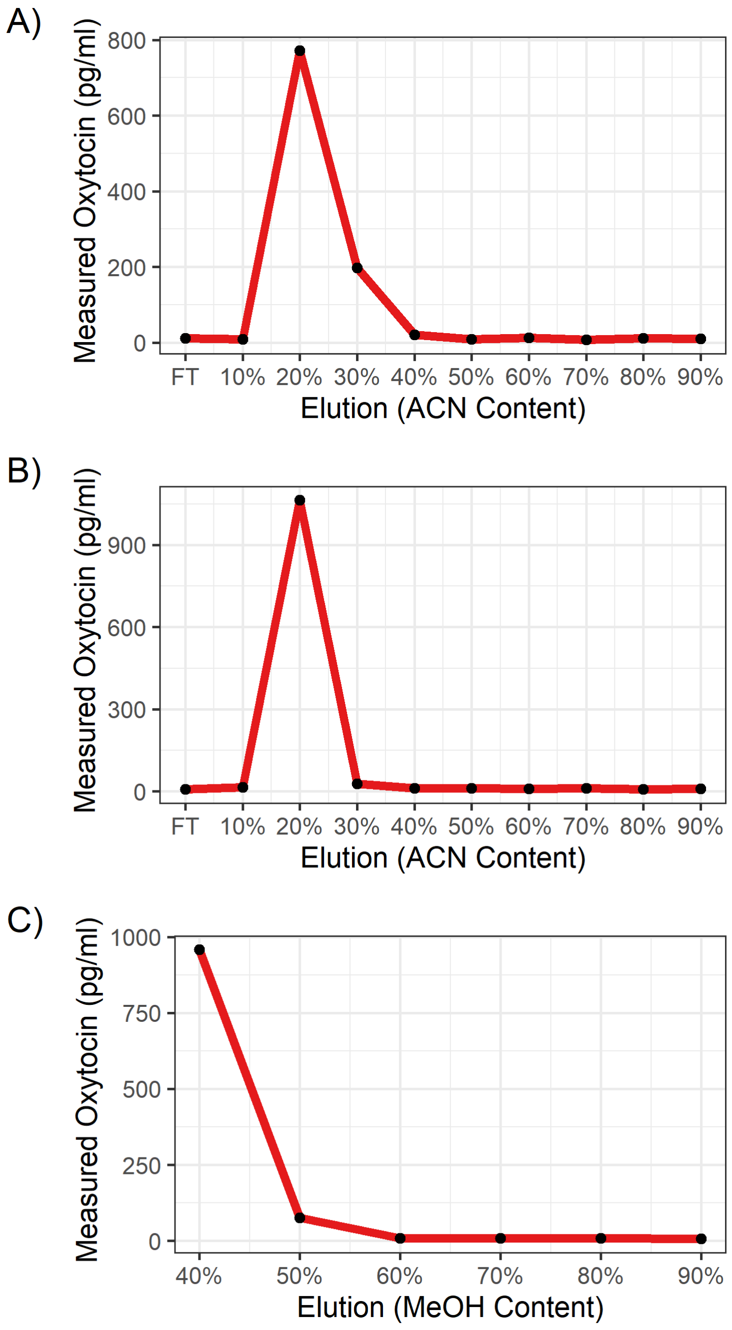

**Figure A1:** Sequential elutions with increasing organic content on a single cartridge of each type. A sample of oxytocin in assay buffer (approx. 1,000 pg/ml) was loaded on each cartridge. FT = "Flow Through" (from the sample loading). A) HLB: No measurable oxytocin eluted at 10%, the majority eluted at 20%, and some additional oxytocin eluted at 30%. B) StrataX: almost all of the sample elutes at 20% ACN. C) MCX: most of the sample elutes at 40% MeOH, with the remaining oxytocin eluting at 50%.

oxytocin (but hopefully not interfering molecules). To do this, we began with three different elution buffers at pH 7, 9, and 11. All elution buffers were composed of 50% MeOH. The other 50% varied for each buffer: the pH 7 buffer was 50% $H_2O$; the pH 9 buffer was 50% a mix of 20 mM $NH_4HCO_2$ and $NH_4OH$ aqueous solution; the pH 11 buffer was 50% $NH_4OH$ in aqueous solution. We found that oxytocin did not elute at pH 7, eluted slowly at pH 9 (spread over two 1 ml elutions, and did not completely elute), and eluted more quickly at pH 11 (mostly in first 1-ml elution, with some remaining in the second 1-ml elution). Even with pH 11, however, we had only about 70% recovery of our loaded spiked buffer sample. We therefore proceeded with 50% MeOH elution at either pH 11 or higher (details on these experiments below).

##### 3.4 Elution Volume

We tested the effect of elution volume for both cartridges, and found that while more aggressive elution buffers eluted most of the sample in the first 1-ml elution, a second elution step often included some additional sample. The extra elution was most important for less aggressive elution buffers (e.g. lower pH for the MCX cartridge). We therefore increased the elution volume in all cases to 2 ml, split across two 1-ml elution steps.

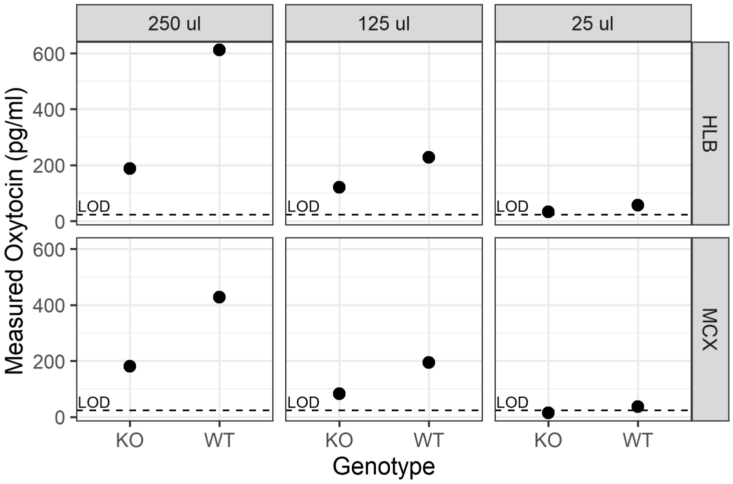

**Figure A2:** Genotype contrast on single pools of each genotype, extracted on either the HLB or MCX cartridges, using three different volumes of sample. While the wildtype pool measures higher than the knockout pool (an improvement from unextracted samples), there is still considerable interference in the knockout samples measured with 250 µl and 125 µl. With only 25 µl of sample, both genotypes measure very close to the limit of detection (LOD, dotted line). It should also be noted that while 250 µl produced a stronger genotype contrast (i.e. higher ratio of WT to KO), there is rarely enough sample volume from an individual collection of a single animal for this to be feasible for analyses of individual differences or specific time points.

#### 3.5 Genotype Contrast

Having determined partially optimized protocols using spiked buffer, we then tested the performance of both methods by again using wildtype and knockout mouse urine, measured on the Arbor Assays kit. We found that extraction improved the genotype contrast, but there was still significant interference in both cases (Figure A2). We also tested using different amounts of urine (thereby measuring at different dilutions post-extraction), and while increased dilution (decreased sample volume) did result in less interference in the knockout samples, there was no point at which the knockout samples were below the lower limit of detection while wildtype samples were sufficiently above the limit of detection to effectively assay their variation.

#### 3.6 Further Extraction Refinement

We therefore attempted to refine both methods over several assays. For the HLB method, the only variable that could be manipulated was the quantity of washing (potentially balanced by a less aggressive wash, but more aggressive was not possible without losing oxytocin). We found that increased wash volumes resulted in some minimal loss of oxytocin in spiked buffer, although increased washing at both 5% and 10% ACN did not improve the genotype contrast, with knockout samples still measuring well above the limit of detection.

It should be noted that the genotype contrast was much better on the Enzo kit (ratio WT/KO = 6.4 - 10.3) than the Arbor kit (ratio WT/KO = 3.4 - 6.3), however the knockout sample pool still measured just above that assay's sensitivity (sensitivity = 15.0 pg/ml; KO pool across conditions = 19.8 - 23.8). The MCX method (after some of the refinements outlined below) performed better on both the Arbor and Enzo kits, and we proceeded primarily with developing the MCX method further.

For the MCX method, we adjusted 1) the pH of loading, 2) the pH, organic content, and quantity of washes, as well as sequential combinations of different washes, and 3) the pH and organic content of the elution. Ultimately, all three steps were refined.

By comparing spiked buffer and dog urine, we found that when loading at a higher pH (tested 2, 3, 4, and 5), oxytocin was still retained, but some signal in the urine was lost, and this loss increased at higher pH (see figure A3). This suggests that there are molecules in urine that interfere with the immunoassay but are not retained on the cartridge when loaded at pH 5. We tested the effect of loading at pH 5 on wildtype and knockout mouse urine, as well as spiked samples of both genotypes, with a sequential elution of pH 10, 11, and 12. We found a negligible genotype contrast in the pH 10 elution (WT = 117 pg/ml; KO = 87 pg/ml), with only some of the exogenous oxytocin eluting from spiked samples. In the pH 11 elution, we found a much stronger genotype contrast (WT = 194 pg/ml; KO = 47 pg/ml). In the pH 12 elution, most of the endogenous oxytocin had eluted from the unaltered samples (WT = 31 pg/ml; KO = 22 pg/ml), although some residual exogenous oxytocin in the spiked samples was still present (approx. 100 pg/ml). From this we concluded that loading at pH 5 was helpful but not sufficient, and that we needed to further refine the washes. (This loading experiment was done without alteration to the wash buffer per se, except for washing at pH 5 because the wash buffer is a combination of the loading buffer and MeOH.)

For the wash step, we found that there was a balance between washing aggressively enough to eliminate interference and washing too aggressively, such that too much oxytocin was lost. More washing, at higher pH, and with higher organic content all washed away both interference and oxytocin, so we needed to find a balance that minimized interference while maximizing oxytocin retention. Not all of this data is shown here, but figure A4 shows the effects of increasing MeOH content (all using a 1 ml $H_2O$, 3 ml wash buffer, 1 ml of $H_2O$ wash, loaded at pH 5, and eluted at pH 11). From this and subsequent experimentation, we settled on a 40% MeOH, 60% loading buffer (pH 5) wash buffer. We then experimented with different wash volumes (figure A5), and concluded that 4 ml was sufficient to wash away most interference, without also washing away too much oxytocin. We therefore proceeded with 4 ml of 40% MeOH, 60% loading buffer (pH 5) wash buffer (preceded and succeeded by 1 ml $H_2O$ washes).

Due to somewhat low recoveries of exogenous oxytocin in a few assays, we increased the pH of the elution buffer to 12, with the hopes of increasing recovery (we then dialed this back to pH 11 later, which minimized interference, as discussed in the main results).

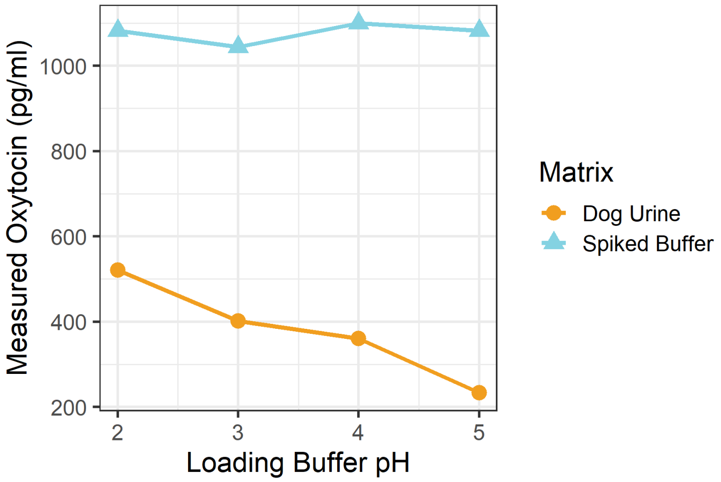

**Figure A3:** Effect of MCX loading buffer pH on retention. Loading at higher pH does not affect the retention of the spiked buffer, but does lead to decreased measurement in dog urine, which may represent the removal of interfering molecules that are no longer bound on the cartridge during loading.

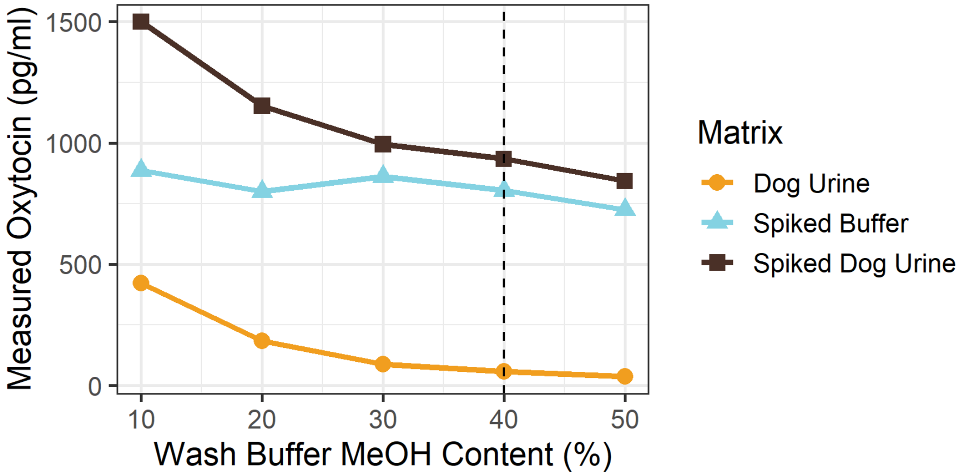

**Figure A4:** Effect of MCX wash buffer organic (MeOH) content on retention and interference. Washes of 10-30% MeOH seem to remove interference and not oxytocin. Washing with 50% MeOH appears to be too aggressive, beginning to wash away more of the oxytocin itself (seen in the spiked buffer). We therefore settled on a 40% MeOH wash buffer. Assay #2355.

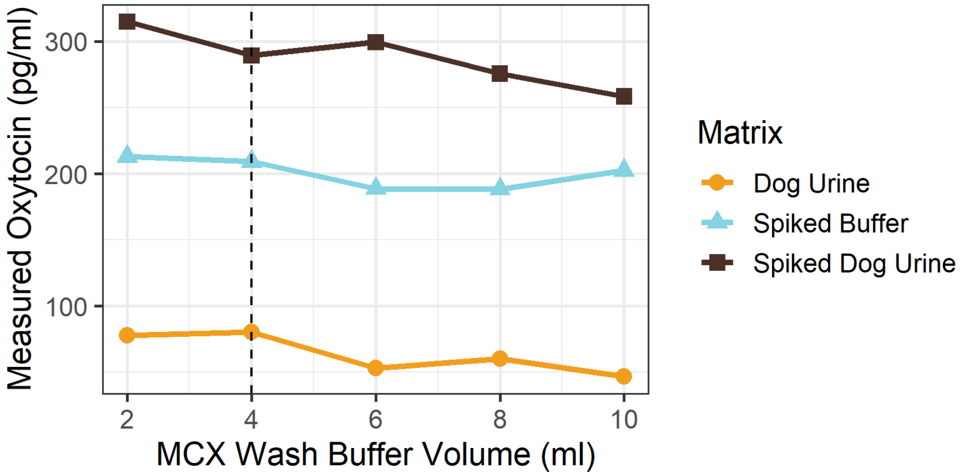

**Figure A5:** Effect of MCX wash buffer volume on retention and interference, all using 40% MeOH. Increased washing generally results in decreased measurement. There was a particularly noticeable decrease in both the spiked buffer and dog urine samples at 6 ml. We therefore proceeded with a 4 ml wash.

#### 4 Epitope Mapping

One set of Pepscan fine-mapping results are shown in figure A6. The complete raw data from the epitope mapping is provided in a separate file.

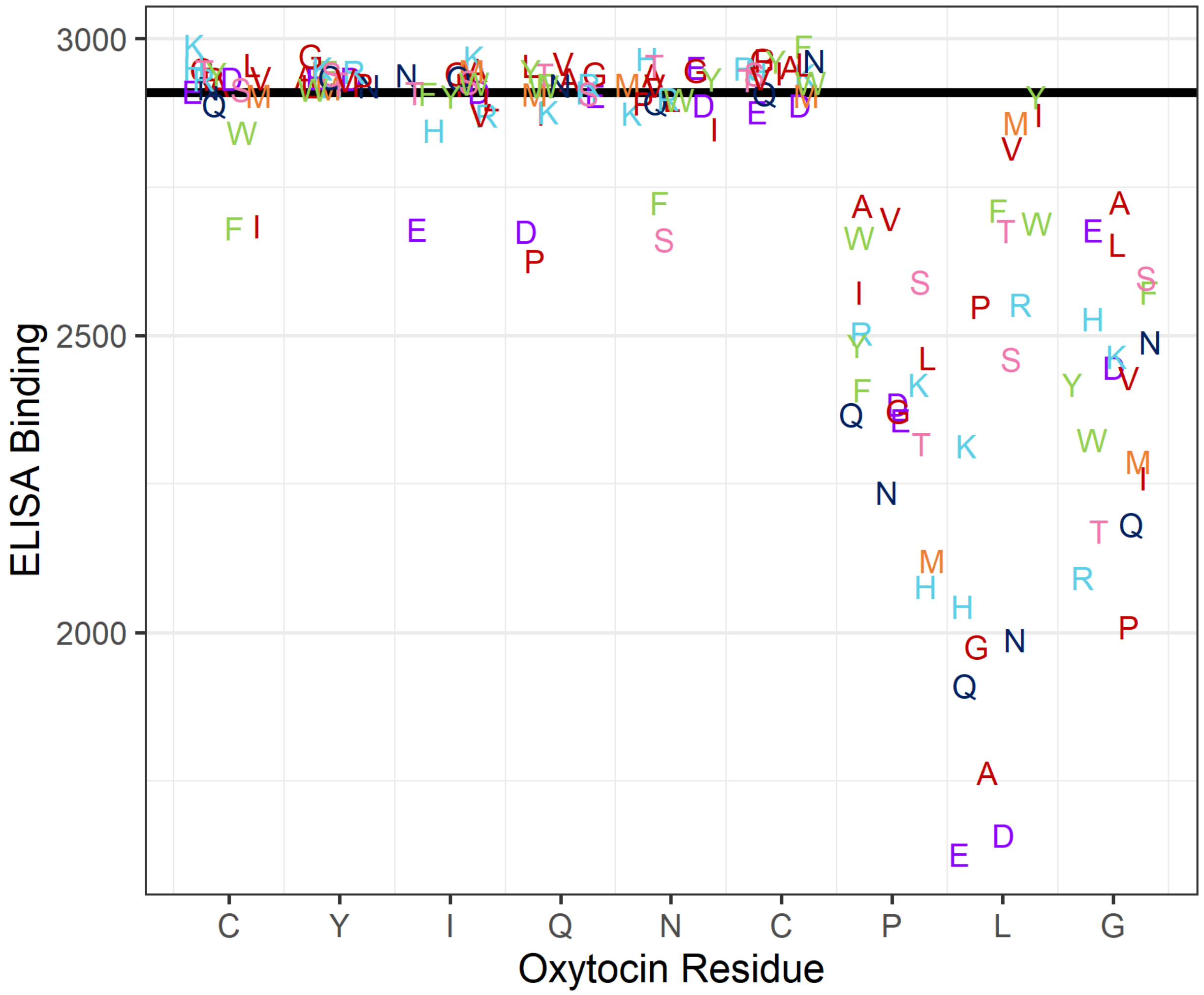

**Figure A6:** Pepscan fine mapping of the core epitope. Starting with oxytocin sequence (x-axis), peptides were synthesized with substitutions at each residue. The black line corresponds to the average binding of oxytocin itself (7 replicates). Peptides with a single substitution are plotted along the x-axis at the residue where the substitution was made (jitter added to minimize overplotting). The one-letter amino acid code is used to indicate the identity of the substitution. The data presented here are for a linear peptide sequence with the antibody at a 1000-fold dilution, but other dilutions and cyclic peptides were also tested (see raw data file).

#### 5 Oxytocin Fragment Cross-Reactivities

Cross-reactivities can be calculated either by weight or by molarity. While figure 5 in the main text shows cross-reactivity-by-molarity, figure A7 below shows cross-reactivity-by-weight. The raw data is given in table A5.

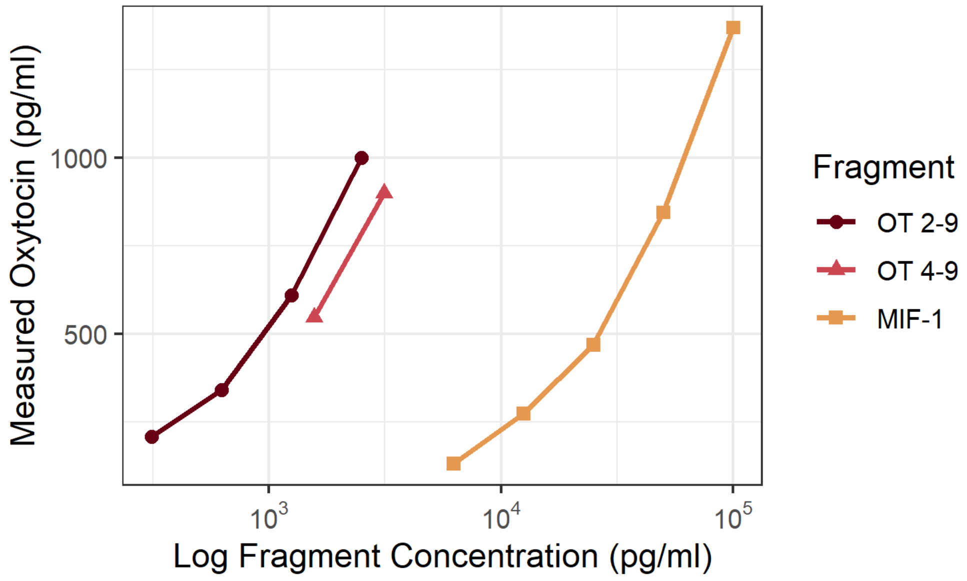

**Figure A7:** Measured oxytocin molar concentrations on the Arbor Assays EIA kit vs. known molar concentrations of three biologically active fragments of oxytocin (log scale). Only measurements within the range of 30-70% of the assay's maximum binding are shown and used to calculate cross-reactivity, the percent of the known fragment concentration that is measured as oxytocin, by molarity. OT 2-9, the fragment most similar to oxytocin, had the highest cross-reactivity (52% by weight, 47% by molarity), followed by OT 4-9 (32% by weight, 21% by molarity). MIF-1, the smallest fragment, displayed the lowest cross-reactivity (2% by weight, 0.6% by molarity).

#### 6 Parallelism Plots

Serial dilutions were conducted on sample pools from all four species to assess parallelism and determine appropriate dilutions for spike recovery analyses and future assays. The mouse and dog samples diluted parallel to the standard curve throughout the range (figures A8 and A10). Sifaka samples were less parallel at high concentrations (figure A9), while human samples were less parallel at low concentrations (figure A11).

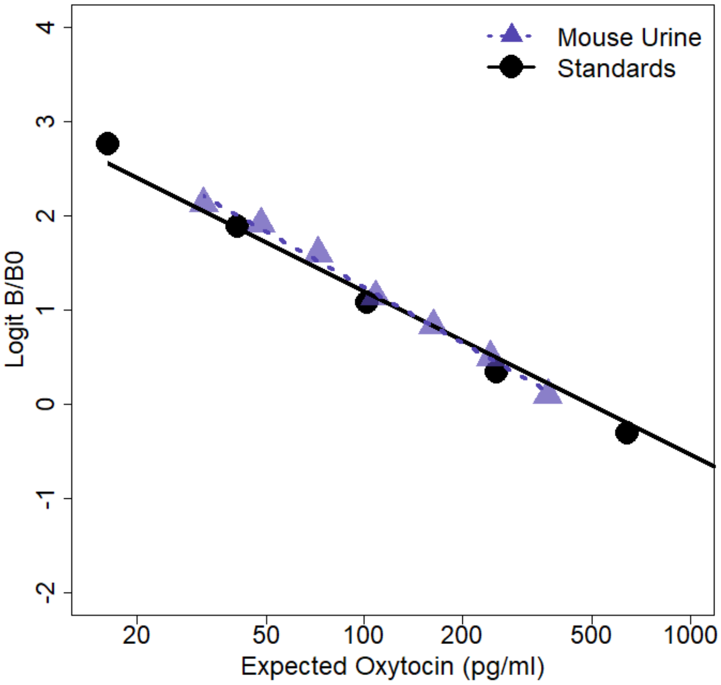

**Figure A8:** Parallelism for a wildtype mouse urine pool, which exhibited good parallelism (CV of corrected concentrations = 10.1%). The first diution (rightmost point) represents a 1.33x concentration (1 ml sample extracted, reconstituted in 750 µl), followed by a 2:3 (p:w) dilution series.

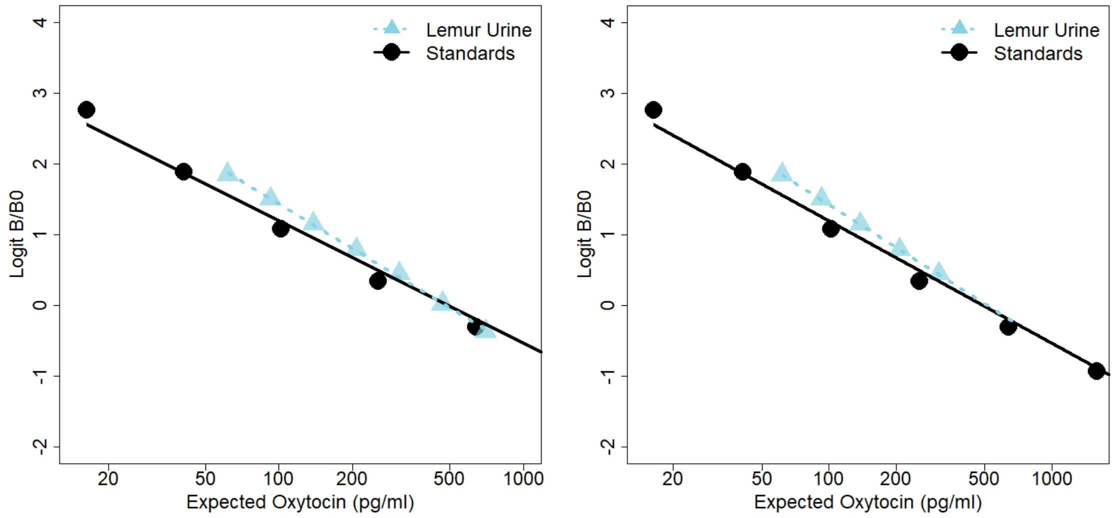

**(a)** Entire Range **(b)** Lower 5 Dilutions

**Figure A9:** Parallelism for a *P. verreauxi* urine pool, demonstrating relatively good parallelism, especially for the lower 5 dilutions. a) Entire range: CV of corrected concentrations = 17.5%. The first diution (rightmost point) represents a 1.33x concentration (1 ml sample extracted, reconstituted in 750 µl), followed by a 2:3 (p:w) dilution series. b) For the lower 5 dilutions, the parallelism improves, with the CV of corrected concentrations dropping to 5.11%.

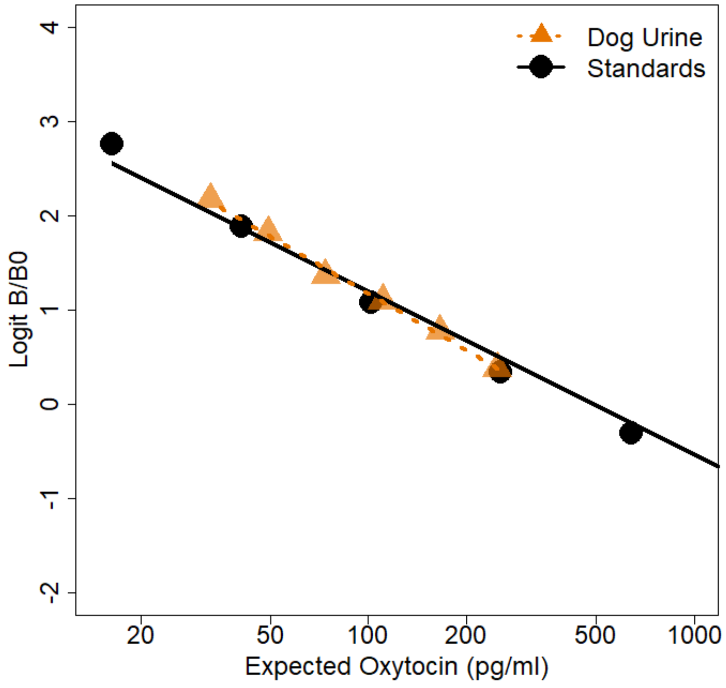

**Figure A10:** Parallelism for a dog urine pool. The parallelism was excellent, with a CV of corrected concentrations of 6.0% across the entire range. The first diution (rightmost point) represents a 5.3x concentration (two 2-ml samples extracted, reconstituted in 750 µl total), followed by a 2:3 (p:w) dilution series.

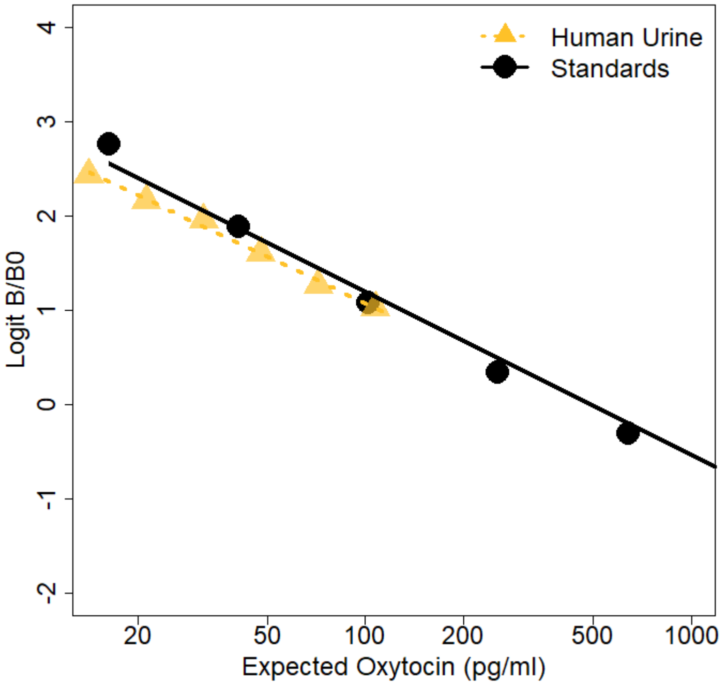

**Figure A11:** Parallelism for a human urine pool. The first diution (rightmost point) represents a 16x concentration (three 4-ml samples, reconstituted in 750 µl total), followed by a 2:3 (p:w) dilution series. The CV of corrected concentrations over the entire range (16x-2.1x) was 18%; excluding the fourth dilution, which had duplicate CV of 25%, this rose to 19%. The CV of corrected concentrations for the first three dilutions was 6.6%.

### 7 Data and Sample Information

#### 7.1 Mice

**Table A2:** Summary of mouse samples included in this study. Experiments 1 and 3 used individual samples, while experiment 2 and all the of the development work was conducted on larger quantities of samples, pooled across wildtype and knockout individuals. Multiple different pools were used throughout methods development. F = female; M = male.

| Description | Section | Strains | F | M | # Across Pools |
|---|---|---|---|---|---|
| Unextracted WT Parallelism | Appendix | B6 | | 1 | |
| Experiment 1: Unextracted Urine | 3.1 | Oxt +/+ | 7 | 3 | |
| | | Oxt -/- | 8 | 6 | |
| Experiment 3: Individual Samples | 3.3 | Oxt +/+ | | 6 | |
| | | Oxt -/- | | 10 | |
| Experiment 2 & Development Assays | 3.2, Appendix | WT (Oxt +/+ & B6) | | | 76 |
| | | Oxt -/- | | | 62 |

**Table A3:** Raw data for the experiment 3 genotype comparison. Corrected concentrations are calculated by multiplying the concentration factor inherent in the extraction (100 µl sample used, reconstituted in 250 µl assay buffer) and are given only for those samples that measured above the assay's limit of detection (22.9 pg/ml). **Abbreviations:** Indiv= Individuals pooled to form single sample. Meas. OT = Measured Oxytocin. Conc. = Concentration. Corr. OT = Corrected Oxytocin.

| # | Indiv | Code | Genotype | Meas. OT (pg/ml) | CV (%) | Conc. Factor | Corr. OT (pg/ml) |
|---|---|---|---|---|---|---|---|
| 1 | 1 | A1 | KO | 15.972 | 3.65 | 2.5 | |
| 2 | 1 | A2 | WT | 26.737 | 14.99 | 2.5 | 66.843 |
| 3 | 1 | A1 | KO | 16.804 | 7.28 | 2.5 | |
| 4 | 1 | A2 | WT | 90.904 | 14.21 | 2.5 | 227.260 |
| 5 | 3 | A1 | KO | 15.738 | 7.45 | 2.5 | |
| 6 | 1 | A2 | WT | 71.331 | 0.83 | 2.5 | 178.328 |
| 7 | 2 | B1 | KO | 16.511 | 22.60 | 2.5 | |
| 8 | 1 | B2 | WT | 85.476 | 5.65 | 2.5 | 213.690 |
| 9 | 1 | B1 | KO | 7.582 | 0.27 | 2.5 | |
| 10 | 1 | B2 | WT | 57.577 | 1.67 | 2.5 | 143.943 |
| 11 | 2 | B1 | KO | 7.431 | 57.17 | 2.5 | |
| 12 | 1 | B2 | WT | 128.688 | 7.29 | 2.5 | 321.720 |

#### 7.2 Sifakas

**Table A4:** Individual urine samples used in pools for validation analyses. All of the samples collected in 2019 were flagged as having been potentially mixed with other individuals, and thus targeted for these purposes.

| Sample ID | Name | Collection Date | Sex |
|---|---|---|---|
| II-2 | Unknown | 6/23/2017 | Unknown |
| XI-2 | Albert | 6/23/2017 | Male |
| U044 | William II | 1/28/2019 | Male |
| U075 | Valdes IV | 2/4/2019 | Male |
| U100 | William II | 2/12/2019 | Male |
| U136 | Mafia VI | 2/18/2019 | Male |
| U209 | William II | 3/10/2019 | Male |

##### 7.3 Cross-reactivity Data

**Table A5:** Cross-reactivity data for three biologically active fragments of oxytocin, calculated both by-weight and by-molarity. **Abbreviations:** Meas. OT = Measured Oxytocin, CR = Cross-Reactivity.

| | | By-Weight | | | By-Molarity | | |
|---|---|---|---|---|---|---|---|
| **Fragment** | **Binding (%)** | **Input (pg/ml)** | **Meas. OT (pg/ml)** | **CR (%)** | **Input (pM)** | **Meas. OT (pM)** | **CR (%)** |
| OT 2-9 | 34.5875 | 2500 | 1000.80 | 40.03 | 2482.15 | 993.66 | 40.03 |
| OT 2-9 | 42.6935 | 1250 | 608.93 | 48.71 | 1241.08 | 604.58 | 48.71 |
| OT 2-9 | 53.1165 | 625 | 339.58 | 54.33 | 620.54 | 337.16 | 54.33 |
| OT 2-9 | 62.0295 | 312.5 | 207.52 | 66.41 | 310.27 | 206.04 | 66.41 |
| OT 4-9 | 36.2305 | 3125 | 900.48 | 28.82 | 4497.54 | 894.05 | 19.88 |
| OT 4-9 | 44.598 | 1562.5 | 546.30 | 34.96 | 2248.77 | 542.40 | 24.12 |
| MIF-1 | 30.044 | 100000 | 1370.09 | 1.37 | 351666.90 | 1360.31 | 0.39 |
| MIF-1 | 37.2315 | 50000 | 844.96 | 1.69 | 175833.45 | 838.93 | 0.48 |
| MIF-1 | 47.315 | 25000 | 468.67 | 1.87 | 87916.73 | 465.33 | 0.53 |
| MIF-1 | 57.091 | 12500 | 272.97 | 2.18 | 43958.36 | 271.02 | 0.62 |
| MIF-1 | 69.8825 | 6250 | 131.26 | 2.10 | 21979.18 | 130.32 | 0.59 |
